# Horizontal transfer of ICE*clc*-like elements in *Pseudomonas aeruginosa* clinical isolates

**DOI:** 10.64898/2026.01.08.698369

**Authors:** Valentina Benigno, Nicolas Carraro, Marjorie Gardet, Hanna Budny, Jan Roelof van der Meer

**Affiliations:** Department of Fundamental Microbiology, University of Lausanne, 1015 Lausanne, Switzerland

**Keywords:** integrative and conjugative element, *clc* element, bacterial evolution, *Pseudomonas aeruginosa*, horizontal gene transfer, copper ions

## Abstract

Integrative and conjugative elements (ICEs) are widespread autonomous mobile DNA within bacterial chromosomes. ICEs contain the genes necessary for excision from the chromosome, conjugative transfer to a new recipient cell, and chromosomal reintegration. They can also carry accessory genes that, while not essential for transfer, confer adaptive phenotypes to the host, contributing to host survival under stressful or changing conditions. Genome studies have indicated that *Pseudomonas aeruginosa* clinical isolates carry a wide range of related ICEs with adaptive genes enriched for heavy metal resistance and efflux systems, however, their mobility has remained understudied. Here, we studied the activation and transfer mechanisms of a representative subset of ICE*clc*-type elements. We found that ICE excision could be induced in *P. aeruginosa* by ectopic expression of BisDC, the known master regulator of ICE*clc* activation, pointing to a similar regulatory cascade. A number of elements could be transferred to *P. putida*, where they conferred increased tolerance to specific heavy metals. We also assessed ICE excision rates in response to different classes of stressors using qPCR-based quantification. Sub-lethal copper exposure significantly increased ICE excision rates in several *P. aeruginosa* strains, although this response was strongly strain-dependent and absent in isolates with enhanced copper tolerance, highlighting the importance of host background. Despite elevated excision, copper did not stimulate ICE transfer or induce conjugation gene expression, indicating that ICE excision and conjugation can be uncoupled processes. Transcriptomic analyses revealed strain-specific regulatory responses to copper stress, including differential activation of metal-responsive regulators, oxidative stress pathways, and virulence-associated systems.

**IMPORTANCE:** Integrative and conjugative elements (ICEs) play a major role in bacterial adaptation by mediating horizontal gene transfer, yet the environmental cues governing their activation remain poorly understood. Here, we demonstrate that ICE*clc*-type elements in *Pseudomonas aeruginosa* are transferable at low frequencies and that their excision can be selectively induced by specific stress conditions, notably copper exposure and hypoosmotic stress. Our findings reveal that ICE excision and conjugative transfer can be uncoupled and are strongly influenced by host genetic background, underscoring the complexity of ICE regulation. This work aimed to explore whether clinical conditions or antimicrobial treatment could inadvertently promote ICE-mediated gene transfer, with implications for understanding the evolution of antibiotic resistance and virulence.

## INTRODUCTION

*Pseudomonas aeruginosa* is a highly adaptable opportunistic pathogen that thrives across diverse environmental niches as well as in clinical settings. Its adaptive success is strongly influenced by the extensive variability that exists among strains (1–3). Although most *P. aeruginosa* isolates share a highly conserved core genome, substantial phenotypic diversity arises from differences in accessory gene content, variability in regulatory mechanisms and the presence of mobile genetic elements. Among these, integrative and conjugative elements (ICEs) play a key role, as horizontal gene transfer through ICEs enables rapid acquisition of new traits (4–6). ICEs are mobile genetic elements that integrate into the bacterial chromosome, where they persist during chromosome replication and cell division. In response to specific (but frequently unknown) signals, ICEs activate their excision from the host chromosome and are then thought to prepare for conjugative transfer. Excision results in formation of a circular DNA intermediate, which undergoes nicking, unwinding and is transferred as a single-stranded DNA through a type IV conjugative system to a new host, where it replicates to reform a double-stranded DNA that site-specifically reintegrates into the chromosome (7, 8). ICEs encode all the necessary genes for excision, integration and transfer. In addition to the core genes required for their mobilization and transfer, ICEs often carry “cargo” genes that are non-essential for transfer but can confer adaptive advantages to the host, such as antibiotic or heavy metal resistance, or providing specific catabolic functions (7–10). ICE activation and excision is a rare and tightly regulated event, and a consequence of host exposure to highly specific and often stressful conditions. For example, ICE*clc* in *Pseudomonas knackmussii* is activated in 3-5% of stationary phase cells after growth on 3-chlorobenzoate and transfers after new nutrients are provided to the cells (11, 12). ICE*Bs1* in *Bacillus subtilis* is excised and transferred in presence of a high density of cells lacking ICE*Bs1* and global DNA damage of the host cell (13). ICE*SXT* in *Vibrio cholerae* is induced by the SOS-response (14). On the other hand, CTn915 in *Enterococcus faecalis* is induced by ribosome-targeting antibiotics (15, 16).

A well-characterized family of ICEs is the ICE*clc* family, originally described in *P. knackmussii* B13 and then mostly characterized in *P. putida* (17, 18). ICE*clc*-like elements share a conserved core region encoding the genes responsible for ICE regulation, excision, integration and conjugation, interspersed with variable cargo genes that distinguish each individual element (19–21). The ICE*clc* regulatory cascade for transfer activation is well characterized and several of the regulatory genes are conserved among *clc*-like ICEs (19). The first step in the cascade is considered to be the transcriptional regulator MfsR, which represses its own transcription and that of two other regulatory genes, named *marR* and *tciR* (22). Deletion of *mfsR* results in overexpression of *tciR*, triggering a strong increase in the proportion of cells with activated ICE*clc* (22). TciR activates the regulatory gene *bisR*, and BisR, on its turn, activates the downstream *P_alpA_* promoter, which controls the expression of an operon that includes *bisD* and *bisC*. BisD is a ParB-like protein that slides unidirectionally over the ICE-DNA starting from the *bisS* boxes, and coactivates further ICE*clc* core promoters in association with BisC (23–25).

In a previous study (19), we characterized the distribution and variation of ICEs belonging to the ICE*clc* family among 181 *P. aeruginosa* strains collected from four different hospitals, with most samples originating from the Lausanne University Hospital. Strains were isolated from infected patients, as well as from environmental sources such as sinks and water pipes within healthcare facilities. Most of the *P. aeruginosa* strains carried ICE*s* with a high sequence conservation to the ICE*clc* core region. In contrast, cargo genes were highly variable to ICE*clc* and were notably enriched, among others, for heavy metal resistance genes, and for genes coding for potential multidrug efflux systems and multidrug resistance proteins. A further notable commonality among these ICEs was the presence of highly conserved homologs of *bisD* and *bisC* (>80% sequence conservation with ICE*clc*), but variability in the presence of the ICE*clc* regulators *bisR*, *tciR* and *mfsR. bisR* was present in all characterized *P. aeruginosa* ICEs with >70% nucleotide identity, whereas *tciR* was present in many but not all ICEs, with 70-75% nucleotide identity. Finally, no *mfsR* homologs were identified among the *P. aeruginosa* ICEs. This confirmed a previous observation suggesting that *mfsR* was an evolutionary novelty in ICE*clc* (26) and suggests that the signal(s) that triggers ICE transfer activation in *P. aeruginosa* is distinct from ICE*clc* but potentially converging through a similar regulatory cascade. We speculated that activation may be triggered by heavy metal exposure, or some other type of cell stress or damage.

Our main aim was to test whether the ICE*clc*-type elements in *P. aeruginosa* are transferable and to uncover any conditions that trigger their transfer. To approach this experimentally, we selected a subset of different *P. aeruginosa* ICE-strain combinations, and measured ICE-excision by amplification of the circular DNA junction under growing and stationary phase conditions. We further transformed a number of *P. aeruginosa* with plasmids expressing *bisDC* from ICE*clc* or from the native *P. aeruginosa* ICE, to examine whether ICE activation could be forced through BisDC overexpression. In absence of selectable markers, we attempted to capture transferring and integrating ICEs in *P. putida* with help of a fluorescent trap (27), enabling *a posteriori* testing of ICE-mediated phenotypes, notably, heavy metal tolerance. Finally, we measured ICE excision rates in *P. aeruginosa* by quantitative PCR after exposure to antibiotics or heavy metals, DNA damaging or oxidative stress reagents, osmotic stress, temperature changes and pH variations. Several of the *P. aeruginosa* ICEs were transferable at very low rates, but excision for one ICE was significantly induced in presence of copper and hypoosmotic stress. We analysed the transcriptomic differences in the donor strain in presence or absence of Cu and discuss the potential underlaying mechanisms for this induction.

## RESULTS

### *P. aeruginosa* ICE excision can be forced by overexpression of the known ICE*clc* regulators

A subset of the *P. aeruginosa* clinical isolates were selected to be tested for ICE excision and transfer, based on the variation in the types of *clc-*like ICEs and host strain phylogenetic diversity (Table S1) (19). ICE-excision was measured by PCR amplification for the unique DNA junction formed by circularization of the element in stationary phase cultures. As a proxy for ICE excision rate, we compared qPCR copy numbers of circularized ICEs and of the host chromosome in the samples (Fig. 1A). ICE excision was detected in all but two strains (Fig. 1B; strains 37327 and 19TCP). Most strains exhibited excision frequencies ranging from approximately one event per 1,000 to 100,000 cells. Notably, three strains – 36935, 13520, and 17090 – displayed clearly higher excision rates, reaching about one event per 50–100 cells (Fig. 1B).

**Fig. 1.**
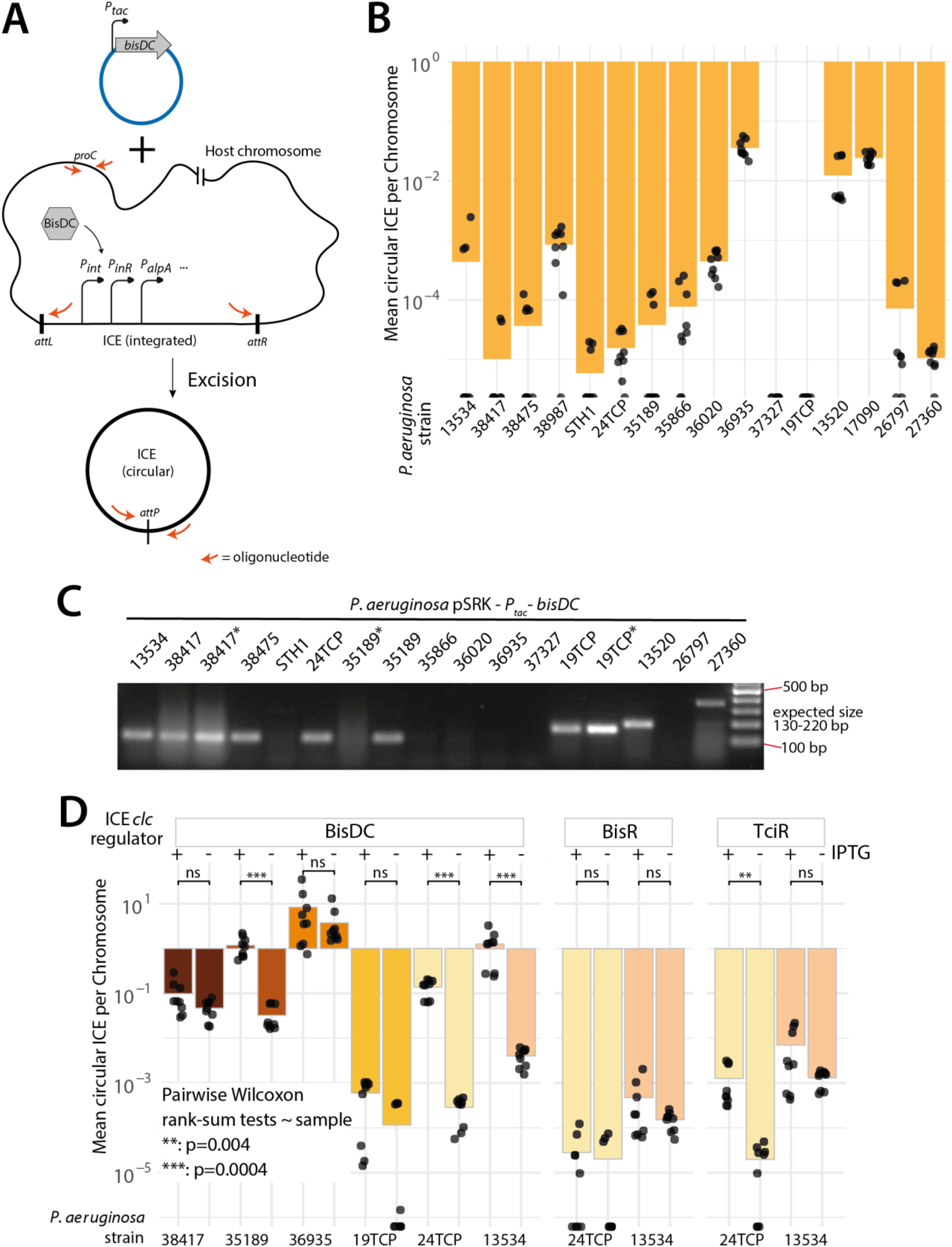
Spontaneous and induced excision of *P. aeruginosa* ICEs by overexpression of ICE*clc* regulators. (A) Schematic overview of the assay used to monitor ICE excision. Upon IPTG induction of the *P_tac_* promoter, BisDC (or another regulator) is overexpressed and activates the ICE core promoters (*P_int_*, *P_inR_*, *P_alpA_*, etc.). No PCR product is formed from integrated ICE but only from excised and covalently closed ICE. (B) Ratio of circular ICE molecules to chromosomal copies, by qPCR on total DNA extracted from stationary phase LB cultures. (C) ICE excision products upon BisDC induction. In all cases, strains express their cognate BisDC homolog, except for samples marked with “*” which carry a BisDC variant from another ICE-type, as indicated. (D) Ratio of circular ICE molecules to chromosomal copies by qPCR, with or without ectopic expression of BisDC, BisR, or TciR from ICE*clc*. The y-axis represents the mean number of circular ICE molecules per chromosome, and the x-axis shows induced versus uninduced controls for each strain and regulator. Statistical significance was assessed using a non-parametric Wilcoxon test comparing each induced condition to its corresponding control.

To test whether ICE excision can be forced by ectopically expressing homologs of the known regulators of ICE*clc* activation (23), we constructed and transformed plasmids into *P. aeruginosa* for IPTG-controlled overexpression of their native BisDC homologs, or of BisDC, BisR or TciR from ICE*clc* (Table S2). After induction of their native BisDC, the circular form of the various ICEs was detected in many, but not all, tested strains (Fig. 1C, Table S3). In strain 27360, the amplicon size was larger than the expected 220 bp, suggesting a possible sequence insertion or rearrangement at the excision junction. For all other strains, the amplicon size matched the expected value, confirming correct ICE circularization. To assess whether different sequence types of *bisDC* are functional to activate and trigger excision of different ICEs, we transformed three of the strains with a plasmid for overexpression of *bisDC* of a different sequence type (Table S2, Fig. 1C strains marked with a “*”). After BisDC induction, the circular form of the ICE was detected by PCR in two out of the three tested strains. Additionally, we transformed six *P. aeruginosa* strains with a plasmid for overexpressing *bisDC, bisR* or *tciR* from ICE*clc* itself (Fig. 1D, Table S1). Overexpression of BisDC, the most conserved regulator, led to a significant increase in ICE excision rate in three strains (35189, 24TCP, and 13534; Fig. 1D). In contrast, strain 19TCP, whose ICE displays a higher divergence from ICE*clc* (both overall and in *bisDC* sequence) did not respond to BisDC induction, consistent with reduced compatibility. Strains 38417, 35189 and 36935 exhibited an unusually high background signal, most likely due to leaky expression of the *bisDC* from the plasmid as the excision was markedly higher than in the wild-type strain (Fig. 1B).

Overexpression of BisR did not induce ICE excision above background levels in either of the tested strains. In contrast, overexpression of TciR significantly increased ICE excision in strain 24TCP, although the magnitude of induction was lower than that observed with BisDC (Fig. 1D). Taken together, these results indicate that ICEs in *P. aeruginosa* remain functional, and that their regulatory networks are at least partially conserved with the ICE*clc* model.

### Transfer of *P. aeruginosa* ICEs to *P. putida* and expression of an associated adaptive phenotype

As ICE excision from the chromosome is essential but not sufficient for transfer, we next sought to test the transferability of ICEs from *P. aeruginosa*. Because the ICEs did not carry clear phenotypically screenable markers (and antibiotic resistance genes could not be successfully introduced into the clinical *P. aeruginosa* strains), we adopted an experimental design described in Ref (27) for ICE*clc* with a fluorescent trap. The trap consists of an extra copy of the ICE*clc* integration site fused to a promoterless *egfp* gene inserted on the *P. putida* UWC1 chromosome. We expected that if, similar to ICE*clc*, the ICEs in *P. aeruginosa* would carry an outward facing constitutive promoter at the junction of the circularized excised form, their integration into the trap would result in this promoter being placed in front of the *egfp* gene, resulting in fluorescent colonies (27). Previously observed sequence conservation of the *att* sequences between *P. aeruginosa* ICEs and ICE*clc* suggested that they may have a similar outward facing promoter (19). Under regular growth conditions with wild-type *P. aeruginosa* donors and the *P. putida* recipient with the fluorescent trap no fluorescent colonies were detected, but donor strains *P. aeruginosa* 38417, 38475, 24TCP, and 13534 with the plasmid overexpressing BisDC produced fluorescent spots on high density mating plates (Fig. 2A). The spots were purified and whole genome sequencing of fluorescent colonies from each of the four transconjugants confirmed insertion of one ICE copy from the *P. aeruginosa* donor into the artificial target in the *P. putida* chromosome. For the remaining tested strains (Table S1), we never observed fluorescent colonies despite overexpression of BisDC. This demonstrated that at least some of the ICEs in the clinical *P. aeruginosa* strains are transferable.

**Fig. 2.**
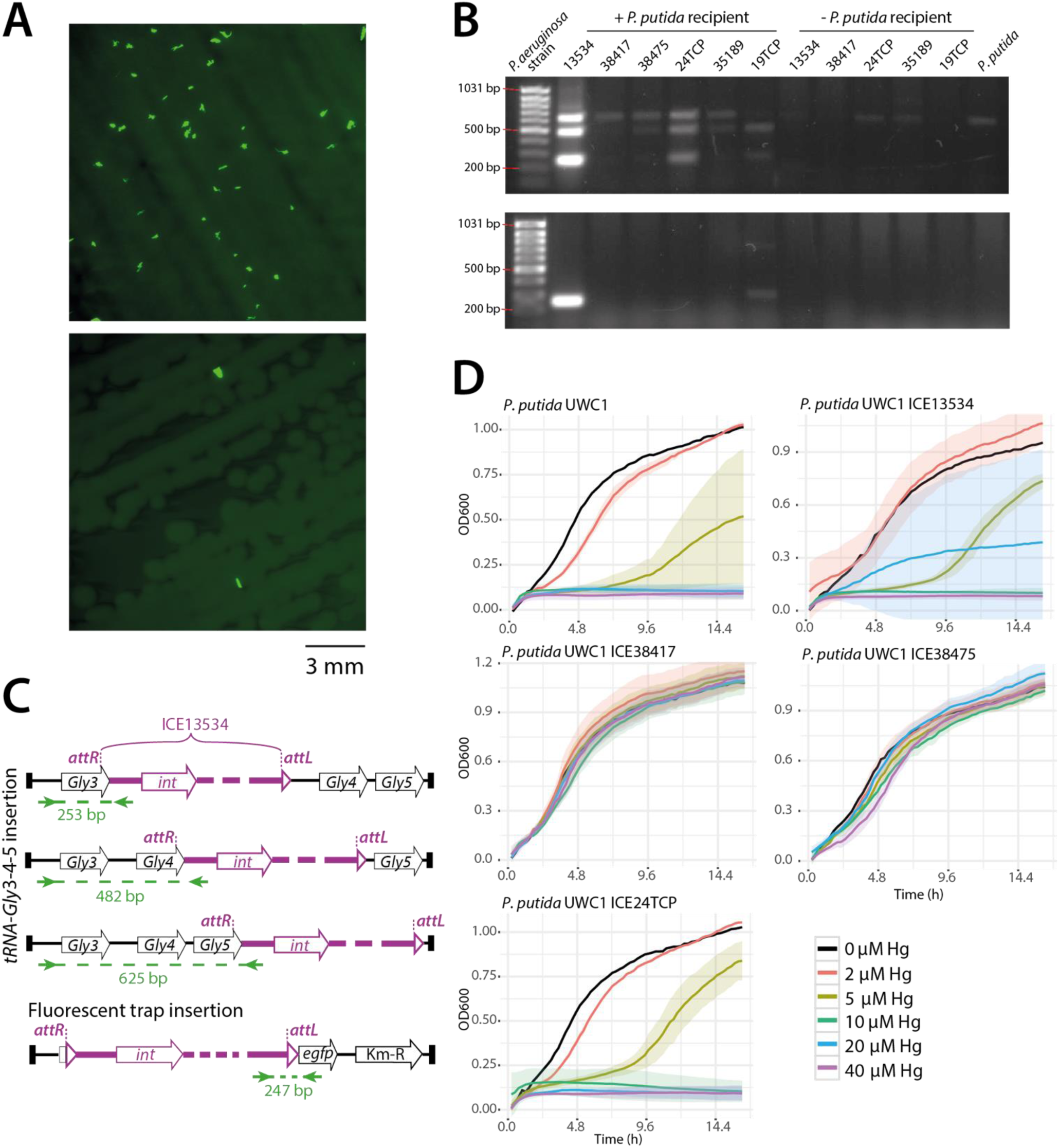
*P. aeruginosa* ICEs transfer to *P. putida* and confer an adaptive phenotype. (A) Mating agarose plates with a coculture of *P. aeruginosa* 13534 and *P. putida tRNA-Gly-gfp*. Highly fluorescent spots correspond to transconjugant colonies. (B) ICE-integration products in transconjugant *P. putida* chromosomes after conjugation with different *P. aeruginosa* strains. DNA bands correspond to amplification of the *tRNA-Gly*3-ICE or ICE-*egfp* junction in *P. putida.* (C) Schematic overview of the amplicon sizes expected from ICE integration into *P. putida tRNA-Gly*3-4-5 or into the fluorescent trap. Example sizes calculated for ICE13534, specific expected sizes vary slightly for different ICEs. (D) Growth curves of *P. putida* parent strain and four transconjugants carrying distinct *P. aeruginosa* ICEs in LB medium with increasing concentrations of mercury.

Because the fluorescent trap is not as efficient as the original insertion site (27) and because the ICEs may insert into four of the other copies of the *tRNA-Gly* genes in the *P. putida* chromosomes, we purified total DNA from the mating mixtures and analysed potential occupation of the insertion sites for the *P. aeruginosa* ICEs by PCR (Fig. 2B-C). We used primers specific for the fluorescent trap and for the *tRNA*-Gly3-4-5 cluster, the most frequent integration sites of ICE*clc* (27) (Fig. 2C, Table S3). Amplification yields varied across strains (Fig. 2B). ICE13534 showed strong amplification for both the native *tRNA*-Gly3-4-5 insertion site and the fluorescent trap, suggesting the most efficient transfer. ICE24TCP only showed integration into the *tRNA-Gly* genes but no detectable product for the fluorescent trap. Other ICEs displayed preferences for specific *tRNA-Gly* loci, e.g., ICE19TCP inserted in loci 3 and 4 but not 5. 19TCP was also the only other ICE to yield a faint band for the fluorescent trap insertion, which was absent in all other strains. These results indicate that integration into native *tRNA-Gly* sites occurs more efficiently than into the artificial target, likely explaining the lack of recoverable transconjugants in some of the attempted *P. aeruginosa* to *P. putida* transfers.

Because of the availability of the recipient strain without the ICE, we could now investigate whether the transferred *P. aeruginosa* ICEs would confer an adaptive phenotype. To better understand the cargo gene content of these ICEs, we analysed their open reading frames (ORFs) beyond conventional sequence-based annotation, which primarily yielded proteins of unknown function (19). We used AlphaFold3 to predict structures of all ORFs lacking functional annotation and compared the resulting models against entries in the Protein Data Bank (PDB). This approach allowed us to assign putative functions to the majority of ICE-encoded ORFs. ICE13534 and ICE24TCP are ∼120 kb long and highly similar, differing by only a few unique ORFs in each element. Both ICEs encode an extensive set of 13 ORFs predicted to be directly involved in copper resistance, as well as a quinolone resistance protein (NorA) (Table S4). ICE38417 and ICE38475 are identical in gene content and contain, among others, operons for mercury and arsenic resistance (Table S4).

Next, we experimentally tested whether *P. putida* carrying the respective ICE would be more tolerant to the heavy metal or quinolone than *P. putida* wild-type. The minimal inhibitory concentration (MIC) for Hg(II) of *P. putida* UWC1 grown in LB was at least 4 times higher for the recipients carrying an ICE from *P. aeruginosa* with detectable Hg-resistance genes (ICE38417 and ICE38475, two identical ICEs coming from strains 38417 and 38475, respectively) than the parental strain (Fig. 2D, Table S5). In contrast, the MICs for As(III) and Cu(II) were unchanged in all ICE-carrying strains compared to the UWC1 parent (Fig. S1A-B, Table S5), likely because the chromosomally encoded baseline resistance of UWC1 is already high and additional ICE-encoded genes do not further enhance tolerance. The MIC for ciprofloxacin did not differ substantially between the transconjugants and the parental *P. putida* UWC1 strain (Table S5), even though *P. aeruginosa* 13534 is classified as ciprofloxacin-resistant (Table S6). This suggests that the resistance determinant present on the ICE may not be functionally expressed or is not contributing to resistance in the *P. putida* background.

### ICE13534 excision is triggered by copper exposure and hypoosmotic stress in *P. aeruginosa* 13534

To test the influence of environmental conditions on ICE excision rates, we chose to focus first on *P. aeruginosa* strain 13534 which belongs to sequence type (ST) 155 (Table S1). This strain carries an ICE, hereafter referred to as ICE13534, that is approximately 118 kbp long and has >80% sequence identity to most of the ICE*clc* core genes. Most of the predicted open reading frames in the cargo region of this ICE have unknown function, but it is also predicted to encode resistance to As and Cu. We focused on a range of conditions that we thought could represent stress and perhaps would lead to increased ICE excision: antibiotics (Table S6), heavy metals (mercury, arsenic, copper), oxidative stress (hydrogen peroxide and paraquat, a herbicide that induces oxidative stress by producing intracellular superoxide radicals (28)), DNA damage (mitomycin C, an intercalating and alkylating agent (29)), osmotic stress (addition of NaCl or media dilution), pH (5 to 9) and temperature stress (shift from 20°C to 45°C). We quantified growth of *P. aeruginosa* 13534 to first identify the conditions and stressor concentrations in which the strain exhibits a measurable growth retardation without complete inhibition. These conditions were then subsequently imposed to quantify ICE excision.

ICE13534 excision in *P. aeruginosa* was quantified by qPCR as the ratio of circular ICE-DNAs and host chromosome in the same sample. As a positive control, we used the strain overexpressing its *bisDC* (Table S1), which we expected would lead to ICE excision in every cell. Indeed, the ratio of circular ICE to chromosome copies in the BisDC-overexpression strain was ∼ 1 (Fig. 3A). As baseline control, we grew the strains in the same media without any specific additional external stress. Under antibiotic exposure, ICE excision was not statistically different than in the baseline control (Fig. 3A), indicating that ICE activation is not induced by the antibiotics and conditions tested here. Similarly, no significant increase in ICE excision was detected after growth with DNA damaging or oxidative stress agents and pH stress, or after temperature shock (Fig. 3A). In contrast, hypoosmotic stress significantly increased ICE13534 excision rates (pairwise Wilcoxon rank-sum test, p = 0.001). No increase was observed after growth in presence of arsenic or mercury (Fig. 3A), but a clear increase of ICE13534 excision was detected after growth in presence of copper, which was statistically significant at all tested concentrations (pairwise Wilcoxon rank-sum test, p < 0.007). Activation under copper stress is optimal at 50 mg/L, and suboptimal at the other tested concentrations (Fig. 3A). This demonstrates that ICE13534 activation is specifically triggered by hypoosmotic stress and copper stress, with a stronger induction at 50 mg/L.

**Fig. 3.**
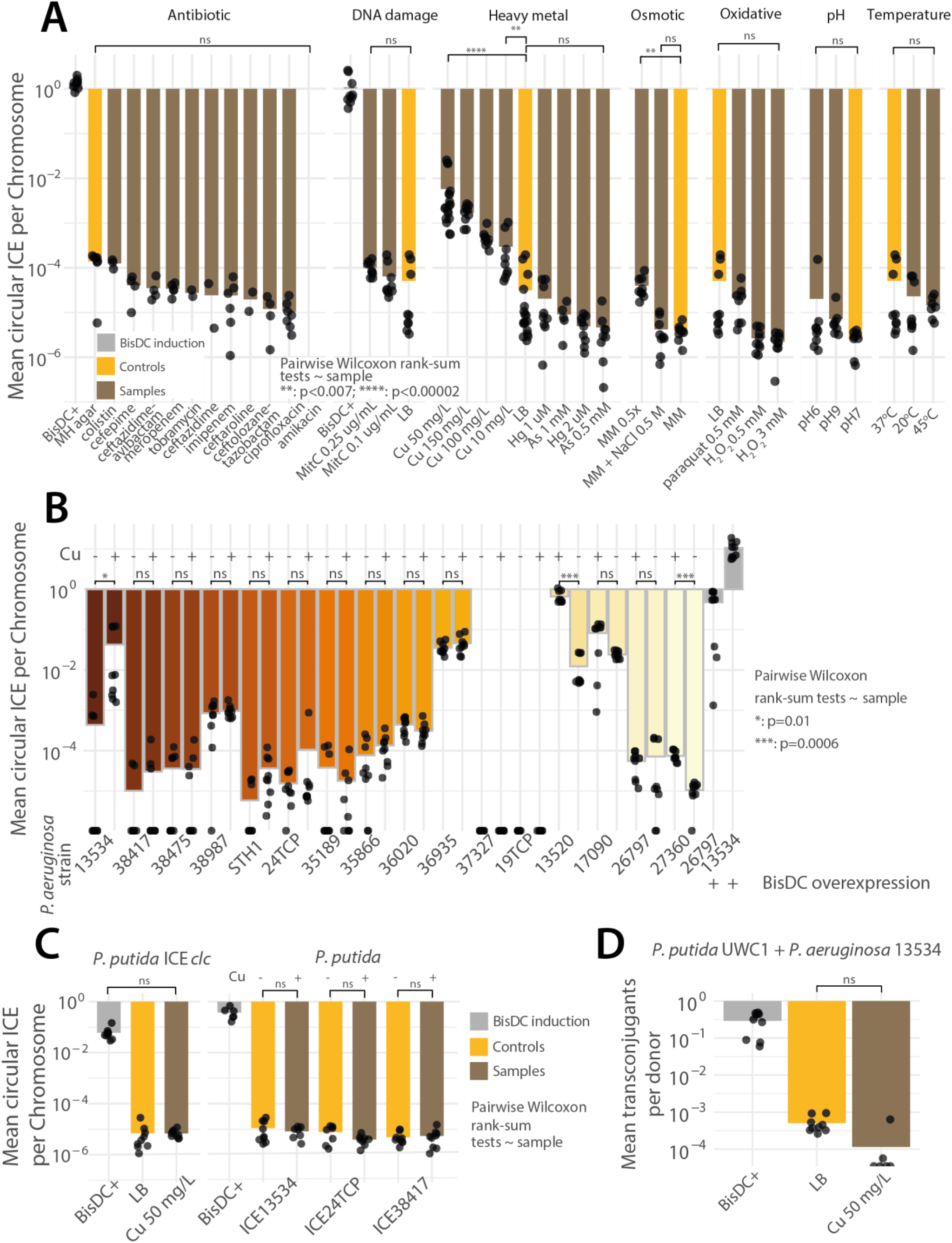
Quantification of ICE excision under stress conditions. (A) qPCR ratios of circular ICE versus chromosome copies of *P. aeruginosa* 13534 exposed to different stresses, as indicated (brown bars), versus control growth (yellow), or strain with BisDC overexpression (grey). (B) As (A), but for ICE*clc*-like elements in other *P. aeruginosa* clinical isolates exposed to copper stress. (C) as (A) but for *P. putida* carrying ICE*clc* or ICEs from *P. aeruginosa* exposed to copper stress. (D) qPCR quantification of *P. putida* transconjugants per *P. aeruginosa* donor, expressed as the ratio of *tRNA-Gly*6-ICE junction amplicons to donor chromosomal copies, in total DNA extracted from mating experiments. Statistical significance was assessed using a non-parametric Wilcoxon test comparing each condition with its corresponding (non-stressed) control.

### ICE13534 activation under copper stress is ICE- and host-specific

We then tested whether ICE13534 excision in response to copper stress was a conserved feature among other *P. aeruginosa* clinical isolates, carrying distinct *clc*-like ICEs. For this, we tested fifteen additional strains (Fig. 3B and Table S1). In most strains, no significant increase in ICE excision rate was observed compared to the non-stressed control (pairwise Wilcoxon rank-sum test, p > 0.05, Fig. 3B). However, copper-induced ICE excision was observed in two ICE–strain combinations, *P. aeruginosa* 13520 and *P. aeruginosa* 27360. Although the ICEs in these strains are not closely related to ICE13534 (86% average nucleotide identity across comparisons), the three hosts are phylogenetically close, and all belong to sequence type ST155 (Table S1). By contrast, ICE24TCP is highly similar to ICE13534 (99% average nucleotide identity), yet its host, *P. aeruginosa* 24TCP, is more distantly related and belongs to a different sequence type (ST164, Table S1). To further evaluate whether the observed copper-induced ICE13534 activation was dependent on the *P. aeruginosa* host, we tested whether ICE13534 activation could be induced in its new *P. putida* host (Table S1), but no significant increase in ICE excision was observed under copper exposure (50 mg/L, Fig. 3C). These findings suggest that host background plays a central role in determining copper-induced ICE excision, potentially more so than the identity of the ICE itself. Finally, we wanted to verify whether copper is also an inducer of ICE*clc* activation. For this, we exposed *P. putida* UWC1 carrying ICE*clc* to copper stress (50 mg/L) and quantified ICE excision rates (Fig. 3C). No significant increase in activation was detected, suggesting that copper-induced activation is not a general property of the ICE*clc*-family of elements but instead depends on specific host–ICE combinations.

Following our findings, we wanted to explore whether ICE13534 copper-induced excision led to increased ICE transfer rates. For this, we performed a conjugation assay with *P. putida* UWC1 (Table S1, not carrying any ICE) as a recipient and *P. aeruginosa* 13534 as a donor. Donor cells were pre-grown in presence of 50 mg/L copper and conjugation mixtures were also incubated in presence of the same Cu-concentration. However, Cu-exposure did not lead to increased transfer rates compared to the baseline (Fig. 3D). These results suggest that, despite significant ICE excision, exposure to Cu does not increase ICE transfer frequency, at least under our experimental conditions.

### Transcriptomic response of *P. aeruginosa* strains 13534, 24TCP, and 13520 to copper stress

To uncover specific host genes that might relay Cu-exposure to ICE excision, we conducted full transcriptome sequencing in *P. aeruginosa* strains 13534, 24TCP, and 13520. ICE13534 and ICE24TCP are nearly identical, but only ICE13534 is excised in response to Cu. In contrast, ICE13520 is shorter (∼80 kb) and distinct in cargo content, lacking predicted metal resistance determinants, but is also excised after Cu-exposure (Fig. 3B). No gene was found to be shared exclusively between ICE13534 and ICE13520. Full genome resequencing suggested 6304, 6047, and 6445 predicted protein-coding genes, respectively, in strains 13534, 24TCP and 13520. Orthogroup analysis identified 5640 orthologous genes shared among all three strains, with a total of 511 strain-specific genes.

Transcriptomes were obtained from 24-h-stationary phase cultures grown in LB with or without 50 mg/L Cu. In presence of copper, strains 13534 and 13520 showed lower yield and decreased growth rate, whereas growth of strain 24TCP was not measurably affected by the used Cu-concentration (Fig. S1C). *P. aeruginosa* 13520 displayed the weakest transcriptional response, with 45 differentially expressed (DE) genes, whereas 13534 and 24TCP exhibited extensive copper-dependent regulation with, respectively, 277 and 257 DE genes (Fig. 4A, Table S7). All three strains have a copy of the known *copRS* and *czcRS* two-component systems involved in metal sensing and resistance, located outside the ICE (30–33). In all strains in response to Cu stress, multiple genes previously reported to be regulated by these Cu-reactive two component systems were DE. This includes genes mediating Cu tolerance, such as *pcoA*, *pcoB*, *copA1,* the operon encoding *ptrA*, the *copI*-homolog PA2807 and *queF* (34). Furthermore, *opdT,* a porin (31), and the heavy metal efflux pump CzcABCD were higher expressed under Cu-exposure (Fig. 4B-D) (35). Multiple pyochelin and phenazine biosynthetic genes were lower expressed with Cu (the latter not present in strain 13520), as expected as a consequence of *czcRS* activation (30). Consistent with long-term copper adaptation (31), the oxidative stress response was not prominent after 24h exposure in strains 13534 and 13520. Strain 24TCP was an exception here, showing significantly higher expressed *ohr*-gene and an overall higher percentage of total transcripts assigned to oxidative stress-related functions in presence of copper (Fig. 4E). The *mexPQ* locus, encoding a general efflux system, was also induced in all strains in presence of Cu. We also found that *catA* and *catC* were strongly upregulated in all three backgrounds under Cu-exposure.

**Fig. 4.**
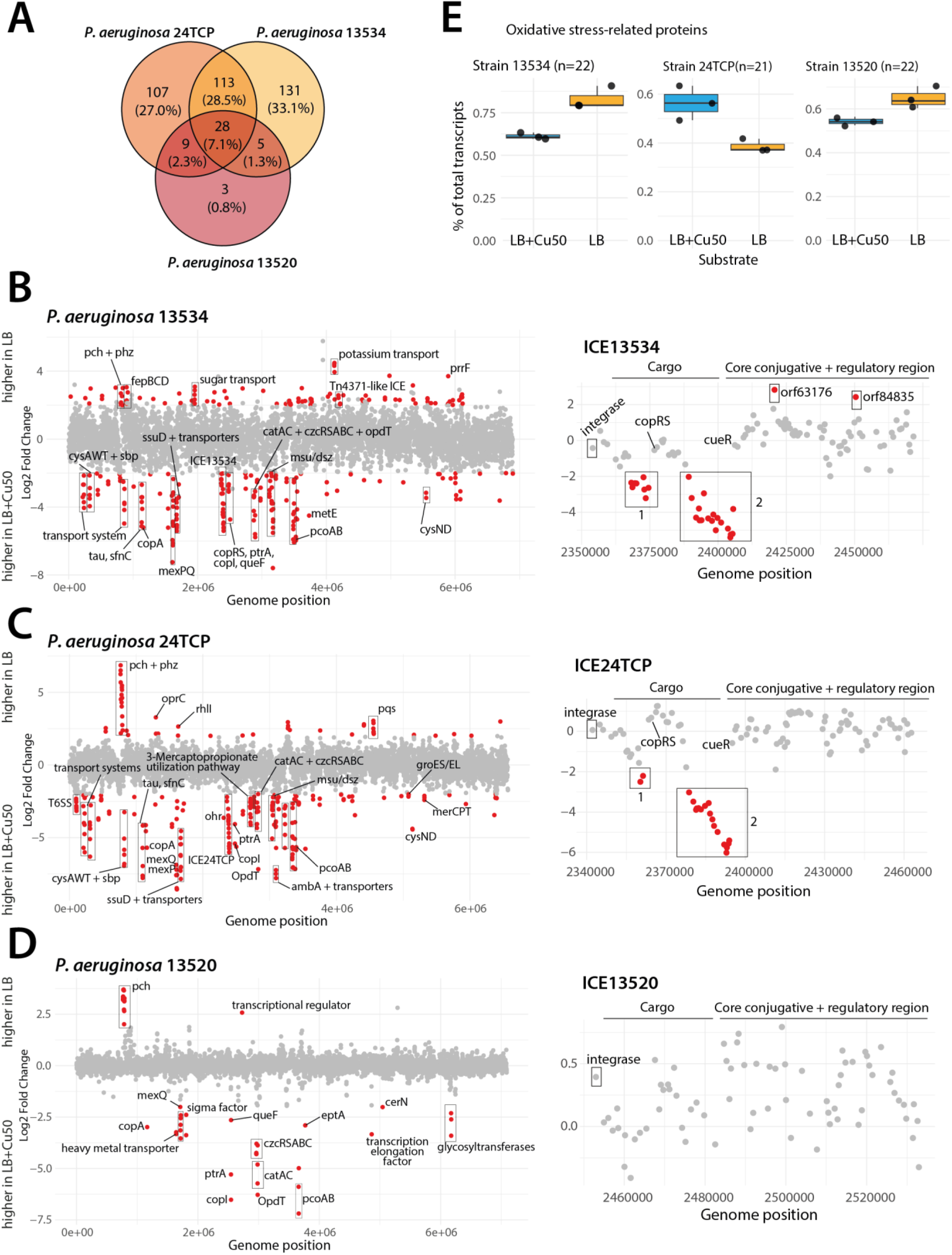
Transcriptomic response of *P. aeruginosa* strains 13534, 24TCP, and 13520 to copper stress. (A) Total numbers of differentially expressed (DE) genes (|log₂ fold change| > 2, *p* adjusted < 0.05). DE genes were calculated separately for each strain by comparing cultures grown in LB versus LB supplemented with 50 mg/L Cu (LB+Cu50). Orthogroup analysis identified DE genes shared among strains or unique to individual strains. (B–D) Distribution of DE genes (red dots) along the chromosome and ICE for each strain. Each dot represents a single gene plotted according to its genomic position and expression value (increased or decreased). (E) Relative transcript abundance of oxidative stress–related genes (n = 21–22 per strain) expressed as the percentage of total transcript counts in LB and LB+Cu50 conditions.

A number of strain-specific transcriptional signatures were also evident (Fig. 4B-D). Cu-exposed strains 13534 and 24TCP showed induction of genes related to sulfate transport and metabolism, indicative of a sulfur starvation response (36, 37). These included *ssuD* and *msuED* for alkanesulfonate desulfonation, and *sfnC, tauD* and taurine transporters for taurine utilization, *dszAC*-family genes for biodesulfurization, the sulfate-regulated *tonB/exbBD* complex, *sbp*, *cysAWTP* for sulfate uptake, and *cysDNHI* for sulfate reduction. Genes uniquely DE in strains 13534 and 13520 included an ECF-type σ⁷⁰ factor, *queF*, and three genes coding for hypothetical proteins of unknown function. Strain 24TCP uniquely upregulated Type VI secretion system (T6SS) genes, and downregulated the quorum-sensing system (*pqsABCDE* and *rhlI* genes) after 24 h exposure to Cu.

No ICE-located genes were DE in strain 13520 (Fig. 4D). In contrast, several cargo genes on ICE13534 and ICE24TCP were strongly upregulated with Cu, whereas genes for the conjugation systems were slightly more expressed in its absence (Fig. 4B-C). Cu-induced ICE gene clusters consisted, first, in a group of seven consecutive genes predicted to be involved in Cu resistance (*pcoA/copABCDKG*), and potentially under control of the ICE-encoded *copRS* two-component system (square “1” in Fig. 4B-C). The level of induction of those genes was higher in strain 13534 than 24TCP. A second cluster comprised 18 genes with mixed predicted functions (Table S4), including *copA* and *norA*. This cluster might be under transcriptional control of a *cueR-*homolog, which is encoded in the opposite orientation adjacent to this region (square “2” in Fig. 4B-C). The terminal gene encoding a glyceraldehyde-3-phosphate dehydrogenase was DE in 13534 but not in 24TCP. In ICE24TCP, an additional insertion of two ORFs (a dehydrogenase and a *tetR*-type regulator, Table S4) was identified immediately upstream of this gene, which may have shielded it from transcriptional readthrough. In summary, the transcriptome of all strains showed an expected Cu-dependent concerted reaction, which for two strains extended into the ICE gene region, but did not show any evident links to induction of ICE excision and/or transfer.

## DISCUSSION

The primary aim of this work was to investigate the mobility of ICE*clc*-family ICEs in *P. aeruginosa*. Indirect evidence from phylogenetic analyses had previously suggested that some of these ICEs were transferred among *P. aeruginosa* strains, and their conserved gene synteny relative to the known transfer modules of ICE*clc* indicated that they are *bona fide* self-transferable elements. To address this, we focused on two levels: (i) quantification of ICE excision and (ii) detection of ICE transfer. In parallel, we examined both wild-type conditions and *P. aeruginosa* strains engineered to ectopically express the known ICE*clc* regulators BisDC, BisR, and TciR, either from ICE*clc* itself or from native homologs encoded on the *P. aeruginosa* ICEs. Finally, we tested whether exposure to diverse stressors could enhance any of the ICE mobility steps.

As expected, wild-type growth conditions in all of the tested *P. aeruginosa* ICEs did not result in any detectable transfer to a *P. putida* recipient, and measured ICE excision mostly remained at frequencies of 10^-4^-10^-6^ per chromosome copy. Ectopic induction of cognate BisDC activators resulted in strongly increased fractions of excised ICE in all tested *P. aeruginosa*. This indicates that the role of BisDC in ICE activation in *P. aeruginosa* is similar as for ICE*clc,* and likely results in similar formation of low proportions of ICE-transfer-competent cells in the population (23). The activation could not in all tested cases be achieved by ectopic expression of the heterologous BisDC regulators from ICE*clc* itself, suggesting that selective differences exist through which cellular or environmental cues are integrated by the ICE to allow its activation. Particularly overexpression of BisR and TciR, which are thought to provide the first level of control on ICE*clc* activation in *P. knackmussii* and *P. putida*, were not effective in inducing ICEs of *P. aeruginosa*. Collectively, these findings would indicate that despite their sequence and structural similarity, the upstream regulators (e.g., TciR and BisR homologs) have become adapted to respond to distinct host contexts (in *P. aeruginosa* as opposed to *P. putida*), but converge to similar output and activation of BisDC. This also suggests that the environmental or physiological cues for ICE activation in *P. aeruginosa* differ from those of ICE*clc*. The latter is activated at least 5 orders of magnitude after growth of the host on 3-chlorobenzoate, although this substrate in itself is not the trigger for activation.

Despite wild-type ICE transfer from *P. aeruginosa* to *P. putida* remaining undetectable, we managed to capture some of the ICEs in *P. putida* through BisDC overexpression in the host and usage of a conditional fluorescent trap (in absence of a strong selectable marker on the ICE itself). By using site-specific PCR, we also demonstrated that the transferred ICEs can target various of the expected *tRNA-Gly* integration sites in the new host. This proved that the ICEs are functionally transferable, provided the proper induction conditions are met. We could also show that a number of the ICEs can provide a detectable phenotype in their new recipient *P. putida*, in form of increased tolerance to mercury exposure.

Different compounds and conditions were tested for potential increase of detectable ICE excision rates in *P. aeruginosa*. Much to our surprise, sub-lethal Cu exposure (50 mg Cu^2+^ per L) significantly increased ICE excision rates in *P. aeruginosa* strains 13534, 13520, and 27360. In contrast, no induction was observed in strain 24TCP (among others), despite its ICE being 99% identical at the nucleotide level to that of 13534, suggesting that host background plays a crucial role in determining the response. Indeed, growth of strain 24TCP in presence of Cu was not impaired compared to that of strains 13534 and 13520. We assume, therefore, that the action of Cu^2+^ is indirect and that, for example, the better defense against Cu^2+^ (like in strain 24TCP) would diminish this indirect effect. Copper ions were previously documented to promote horizontal transfer of plasmids and ICEs (38–40). For example, exposure to sub-inhibitory concentrations of copper significantly enhanced conjugation of the SXT/R391 ICE from *Proteus mirabilis* to *Escherichia coli*. This effect was attributed to copper-induced production of reactive oxygen species (ROS) and increased cell membrane permeability (40, 41). Although copper exposure in these three *P. aeruginosa* strains significantly increased ICE excision rates, it did not measurably enhance ICE transfer to *P. putida*. Consistently, transcriptomic data revealed no induction of the ICE-encoded conjugation genes under copper exposure, suggesting that ICE excision and conjugation can occur as distinct, uncoupled processes rather than as a single coordinated event.

Exposure to Cu^2+^ triggered distinct regulatory and physiological responses among the *P. aeruginosa* isolates. Strain 13520, whose ICE lacks metal-resistance genes, showed minimal transcriptional change under copper stress, suggesting a less responsive or sensitive regulatory network. Unlike ICE13534 and ICE24TCP, ICE13520 does not encode additional copies of the *copRS* and *cueR* regulators, which, once activated, likely influence not only ICE-borne genes but also broader chromosomal transcription, potentially explaining the stronger overall response observed in the other two strains. Consistent with previous reports (31), prolonged copper exposure did not show an oxidative stress signature, except in strain 24TCP, where *ohr* expression remained elevated. The *ohr* gene codes for a thiol-dependent peroxidase that protects bacteria from organic hydroperoxide-induced oxidative stress (42). This strain also uniquely upregulated the T6SS and suppressed quorum-sensing genes, potentially reflecting an altered physiological state under copper exposure compared to the two other studied strains. Curiously, copper exposure induced the *catABC* genes in all strains, which are also activated in hosts carrying ICE*clc* and growing on 3-chlorobenzoate. In addition, strains 13534 and 24TCP showed transcriptional signatures consistent with sulfur limitation. Copper toxicity is known to disrupt Fe–S cluster-containing proteins, thereby increasing the cellular demand for sulfur and iron and potentially triggering sulfur assimilation and starvation responses, as previously shown in *Bacillus subtilis* (43).

Together, our results demonstrate that ICE activation, excision and transfer in *P. aeruginosa* can be governed by a conserved regulatory hierarchy centered on BisDC. Copper exposure acts as an environmental cue that promotes ICE excision in a subset of *P. aeruginosa* strains, specifically those exhibiting growth impairment under copper stress, yet without triggering expression of conjugation functions or yielding a measurable increase in ICE transfer itself. This work shows how ICEs are embedded within the host regulatory and stress-response networks, ultimately determining whether environmental cues translate into their horizontal gene transfer.

## MATERIALS AND METHODS

### Strains and culture conditions

*Escherichia coli* DH5α–λpir cells used for plasmid cloning were cultured at 37°C and 180 rpm rotary shaking in LB medium (44) added with the appropriate antibiotic. The following antibiotics concentrations were used, if necessary: gentamicin (Gm) 10 μg mL^-1^, tetracycline (Tc) 50 μg mL^-1^. *P. aeruginosa* and *P. putida* strains were grown at 37°C and 30°C, respectively, and cultured in LB broth; 21C minimal media (MM) supplemented with 10 mM succinate (45); or M63 MM [0.4 g L^-1^ KH_2_PO_4_, 0.4 g L^-1^ K_2_HPO_4_, 2 g L^-1^ (NH_4_)_2_SO_4_, 7.45 g L^-1^KCl] supplemented with 2 g L^-1^ casein hydrolysate, 20 mM sodium benzoate, 1x Hutner’s (46). Cultures were grown overnight with shaking at 180 rpm. For strains carrying plasmids for inducible *bisDC* expression, 1mM IPTG was added when indicated. Strains used in this study are listed in Table S1.

### DNA manipulations

Fragments for plasmid cloning were PCR amplified with NEB Q5 High-Fidelity DNA Polymerase #M0515 and cloned in one step into the digested plasmid (Vazyme ClonExpress II One Step Cloning #C112). The resulting plasmid was amplified in *E. coli* DH5α–λpir cells, purified (Macherey-Nagel NucleoSpin Plasmid kit for plasmid DNA purification) and sequenced. Colony PCR was done to test for successful plasmid transformation, induction of ICE excision, successful ICE transfer and strain identity (Promega GoTaq Green Master Mix #M712). *P. aeruginosa* strains were transformed by electroporation as previously described (47).

Total DNA was extracted using the NucleoSpin® 96 Tissue kit (Macherey-Nagel) using centrifuge processing according to manufacturer’s instruction with minor modifications: samples were pre-treated by addition of 20 µL of 10x lysis buffer (100mM EDTA, 250mM TrisHCl, 5% Tween20) and 20 µL of 50 mg/mL lysozyme. Samples were incubated for 1 hour at 37°C, then, 25 µL of Proteinase K were added and samples were incubated overnight at 55°C. DNA was eluted in a final volume of 40 µL into PCR plates and stored at −20C°. DNA concentrations were measured using Qubit with High Sensivity (HS) dsDNA assay (Thermo Fisher Scientific).

### Determination of MIC of antibiotics and tolerance to stress compounds

Antibiotic susceptibility testing was performed on *P. aeruginosa* strain 13534 using the Test Strip method (Liofilchem) and following EUCAST recommendations (Table S6). The panel of antibiotics was selected based on common administration to treat *P. aeruginosa* infections at the Lausanne University Hospital. A total of eleven antibiotics were tested: ceftazidime, cefepime, ciprofloxacin, meropenem, imipenem, ceftazidime-avibactam, ceftolozane-tazobactam, ceftaroline, colistin, amikacin, and tobramycin. Several colonies were picked from the non-selective plate and resuspend in a saline solution then vortexed to obtain a homogeneous suspension. The bacterial suspension was adjusted to an OD_600_ of approximately 0.06–0.1. Un-supplemented Mueller-Hinton agar was evenly inoculated with the bacterial suspension. Test Strips were then applied to the surface of the inoculated agar. Plates were incubated at 35°C for 16 to 20 hours before reading the MICs.

The remaining assays were performed as growth kinetics in liquid media. To test the growth dynamics of *P. aeruginosa* strain 13534 in different stress conditions, overnight cultures were 100 times diluted in biological triplicates in LB supplemented with the following concentrations of heavy metals, DNA damaging agents, or oxidative stress inducers: 1, 2, 10, 50 µM Mercury; 1, 3, 10, 15 mM Arsenite; 10, 50, 100 and 150 mg/L Copper; 0.25, 0.5, 1, 10 µg/mL Mitomycin C; 0.2, 0.5, 1, 3 mM hydrogen peroxide or paraquat. Growth under osmotic stress was evaluated by diluting overnight cultures in a high-osmolarity medium (21C MM supplemented with 10 mM succinate and 0.5, 0.7, 1, and 1.5 M NaCl) or a low osmolarity medium (MM supplemented with 10 mM succinate diluted 2- and 4-fold). The growth of *P. aeruginosa* 13534 under pH stress was tested by diluting overnight cultures in M63 supplemented with 50 mM of appropriate buffer [sodium citrate buffer for pH 5; 2-(N-morpholino)ethanesulfonic acid (MES) for pH5.5; 3-(N-morpholino)propanesulfonic acid (MOPS) for pH7; [N-tris(hydroxymethyl)methylamino]-2-hydroxypropanesulfonic acid (TAPSO) for pH7.8, cyclohexyl-2-aminoethanesulfonic acid (CHES) for pH 9]. Finally, growth under thermic stress was assessed by diluting overnight cultures in fresh LB and incubating at 15, 20, 25 °C to assess cold shock response, and at 42, 45 °C to assess heat shock response.

For growth under antibiotic stress, the sub-inhibitory condition was defined as the bacterial biomass growing in contact with the antibiotics zone of inhibition and the test strip. For the other stress conditions, sub-inhibitory concentrations were defined as doses that reduced or impacted bacterial growth without completely inhibiting it. The chosen concentration of the agents induced a range of 30-70% decreases in final OD_600_ compared to untreated control.

### Sample collection for ICE activation assays

To screen for conditions inducing ICE activation, cells were cultured as described before under the following conditions: antibiotics (as listed above, on Mueller-Hinton agar with Test Strips); 1, 2 µM Hg; 0.5, 1 µM As; 10, 50, 100, 150 mg/L Cu; 0.1, 0.25 µg/mL Mitomycin C; 0.5, 3mM H_2_O_2_; 0.5 mM paraquat; 21C MM with 10 mM succinate and 0.5 M NaCl; 4-fold diluted 21C MM; M63 supplemented with 50 mM of buffers at pH 6 and 8.2. After 24h of growth, 5 x 10^6^ cells were harvested in 200 µL PBS for DNA extraction. For the antibiotics stress experiment, cells located near the inhibition zone (just below the MIC) and in contact with the Test Strip were collected.

### qPCR assays

qPCR reactions were set up as follows: per reaction, 5 µL of qPCR SYBR™ Select Master Mix, (Thermo Fisher), 0.4 µL of forward primer (10 µM), 0.4 µL of reverse primer (10 µM), 2.2 µL of nuclease-free water and 2µL of DNA sample. The qPCR reactions were run on a QuantStudio 5 or 7 (Applied Biosystems). Standard curves for quantification of chromosomal genes were generated from serial 10-fold dilutions of the gDNA of the corresponding species (*P. aeruginosa* for *proC*, *P. putida* for *rpoD*). To quantify circularized ICE, standard curves were generated using serial 10-fold dilutions of purified amplicons obtained from primers listed in Table S3, mixed in a 1:1 ratio with gDNA to better approximate real sample conditions.

### Statistical treatment of data and reproducibility

qPCR assays were performed in biological triplicates, and each replicate was quantified in technical triplicates. qPCR data was processed in R and data normality was assessed with Shapiro-Wilk test and rejected for several samples. Statistical comparisons between stress conditions and respective negative controls were performed using the non-parametric one-sided Wilcoxon-rank sum test.

### Absolute quantification of ICE, transconjugant and chromosome copies

Copy numbers were calculated based on the initial known DNA concentration and size of the standards. Linear regression equations were obtained from the cycle thresholds (Ct) values against the logarithms of the known copy number of the standard template DNA and were used to directly calculate copy numbers in samples from their Ct values. Copy numbers in the standards were calculated using this formula:

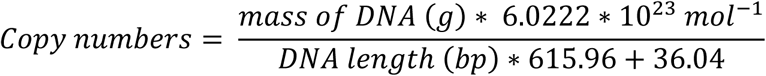

### Conjugation assays

Conjugation assays were performed to assess ICE transfer under wild-type conditions, upon BisDC overexpression, and in response to copper exposure. Donor strains (*P. aeruginosa* ± pSRK-*bisDC*) and the recipient strain (*P. putida* wild-type or carrying an artificial ICE*clc* integration target transcriptionally fused to a promoterless *egfp* gene; Table S1) were grown overnight in LB medium. When relevant, BisDC overexpression in donor cells was induced with 50 mM IPTG for 4 h. For the copper exposure tests, overnight cultures were transferred to fresh LB containing 1mM IPTG and 20 µg/mL Gm (for *P. aeruginosa* 13534 pSRK-Gm-*bisDC38417*) or fresh LB ± 50mg/L copper (for *P. aeruginosa* 13534) and grown for 24h. Subsequently, equal volumes of donor and recipient cultures were mixed and washed in PBS. To isolate fluorescent transconjugants, cell mixtures were spread onto MM agar plates supplemented with 0.5 mM succinate. Plates were incubated at 30 °C for 72 h, and the resulting biomass was subsequently transferred to NA plates for regrowth. Fluorescent biomass was visualized using a Nikon SMZ25 stereomicroscope. For assays testing ICE transfer upon BisDC induction and copper exposure, 20 µL of cell suspension was transferred on top of a 0.2-μm, 13 mm ø cellulose acetate filter (Huberlab, Aesch, Switzerland), that had been placed on a 21C MM agar plate supplemented with 0.5 mM succinate, and 1 mM IPTG or 50mg/L copper according to the preculturing condition. Matings were incubated at 28°C for 72 h, after which cells were recovered from the filters in 1.5 mL PBS by vortexing. DNA was extracted from the resulting cell suspensions and used in PCR assays.

### RNA sequencing

Cells were grown for 24 h in LB ± 50 mg/L Cu from CuSO_4_. Total RNA (>200 nt) was extracted using the NucleoSpin RNA Plus kit (Macherey-Nagel) with on-column genomic-DNA removal. Cells from 20 µL cultures were lysed in 10 mM Tris-EDTA buffer containing 1 mg/mL lysozyme, followed by addition of LBP and BS buffers as per the manufacturer’s protocol. RNA samples were further purified with the RNA Clean & Concentrator-5 kit including in-column DNase I treatment (Zymo Research). Absence of DNA contamination was confirmed by qPCR amplification. RNA-seq libraries were prepared from ∼50 ng input RNA using the Zymo-Seq RiboFree Total RNA Library Kit with a 90-min depletion step and 12 library amplification cycles. Libraries were re-amplified with Q5 Hot Start Polymerase for 12 cycles using Illumina P5/P7 primers and purified by 0.8× CleanNGS bead cleanup (CleanNA). Sequencing was performed on an Element AVITI platform (PE150) at the Lausanne Genomic Technologies Facility, University of Lausanne, Switzerland (https://www.unil.ch/gtf).

### Bioinformatic tools

Complete genomes and ICEs were annotated using Prokka (v1.14.6) (48), guided by the reference proteomes of *P. aeruginosa* PAO1 (NCBI accession number NC_002516), PA14 (NCBI accession number NC_008463), and ICE*clc*. Sequencing reads were quality-trimmed with trimmomatic (v0.39) (49) and aligned to their respective reference genomes using STAR (v 2.7.11a) (50). Samtools (v1.19.2) was used to remove PCR duplicates (51), and gene count tables were generated with HTSeq (v2.0.9) (52). Orthologous genes among strains were identified using OrthoFinder (v3.1.0) (53), and DESeq2 (v1.42.1) (54) was employed for differential expression analysis. AlphaFold3 (55) was used to predict protein structures, which were subsequently compared to the PDB using DALI (56).

## AUTHOR CONTRIBUTIONS

Conceptualization: V.B., N.C., J.R.v.d.M. Methodology: V.B., N.C., J.R.v.d.M. Investigation: V.B., N.C., M.G., H.B. Data curation: V.B., M.G. Formal analysis: V.B., M.G. Software: V.B. Validation: V.B., J.R.v.d.M. Visualization: V.B., J.R.v.d.M. Supervision: V.B., N.C., J.R.v.d.M. Project administration: J.R.v.d.M. Funding acquisition: J.R.v.d.M. Writing—original draft: V.B., J.R.v.d.M. Writing—review and editing: V.B., N.C., M.G., H.B., J.R.v.d.M.

## ACKNOWLEDGEMENTS

This work was supported by the Swiss National Science Foundation grant 310030_204897 to J.R.v.d.M. Funding for open access charge was provided by the Swiss National Science Foundation.

## COMPETING INTERESTS

The authors declare no competing interests.

## DATA AVAILABILITY

The sequencing datasets generated and analysed during the current study are available in the European Nucleotide Archives (ENA) repository under the accession study number PRJEB106170.

## USE OF LLMs

During the preparation of this manuscript, the authors used OpenAI’s ChatGPT to assist with the grammatical refinement of the content. After using this tool, the authors reviewed and edited the content as needed and take full responsibility for the content of the manuscript.

## SUPPLEMENTARY INFORMATION

**Fig. S1.**
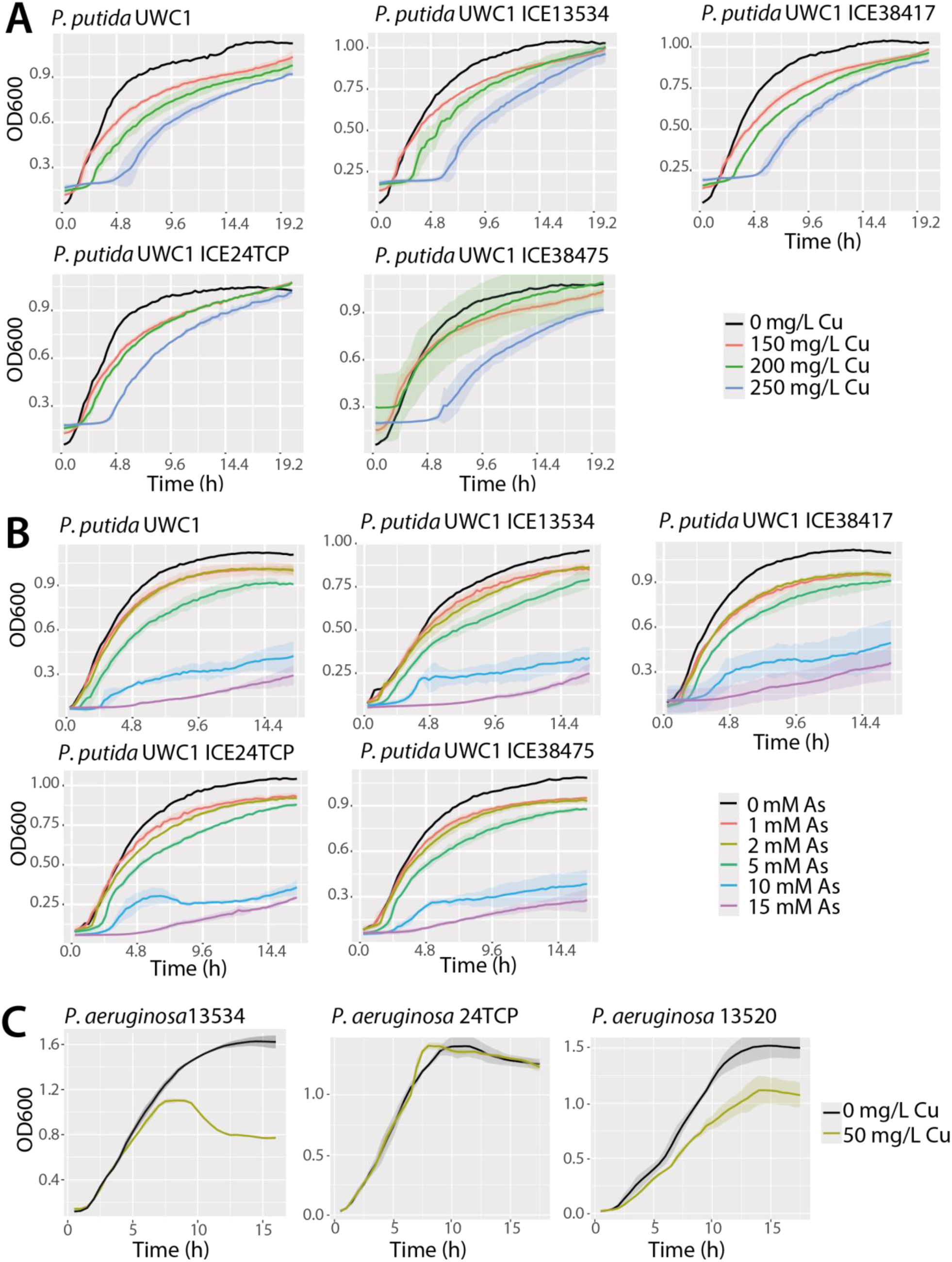
(A-B) Growth curves of *P. putida* parent strain and four transconjugants carrying distinct *P. aeruginosa* ICEs in LB medium with increasing concentrations of copper and arsenite. (C) Growth curves of *P. aeruginosa* 13534, 24TCP, 13520 in LB with or without addition of 50 mg/L copper.

**Supplementary Table 1:**
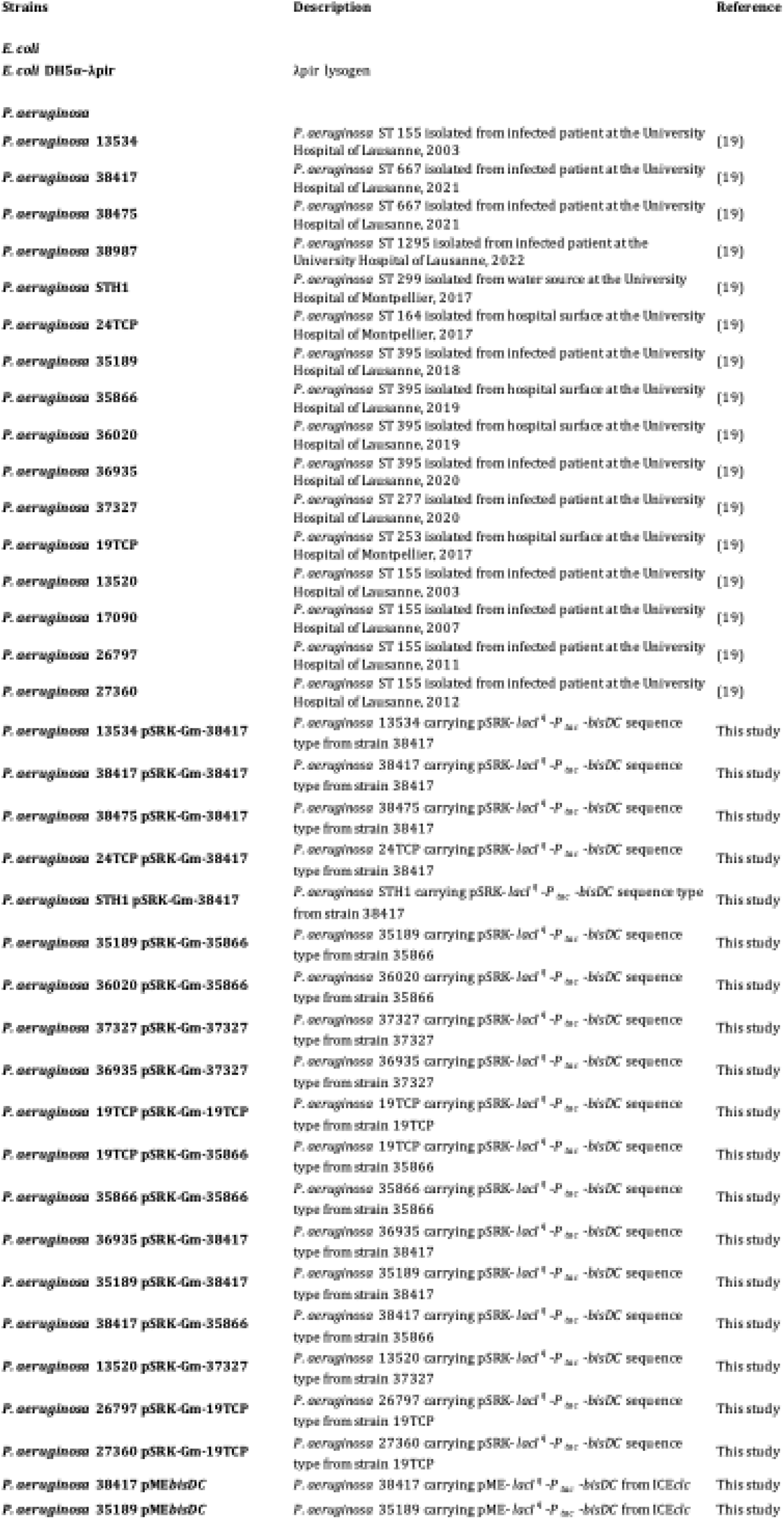
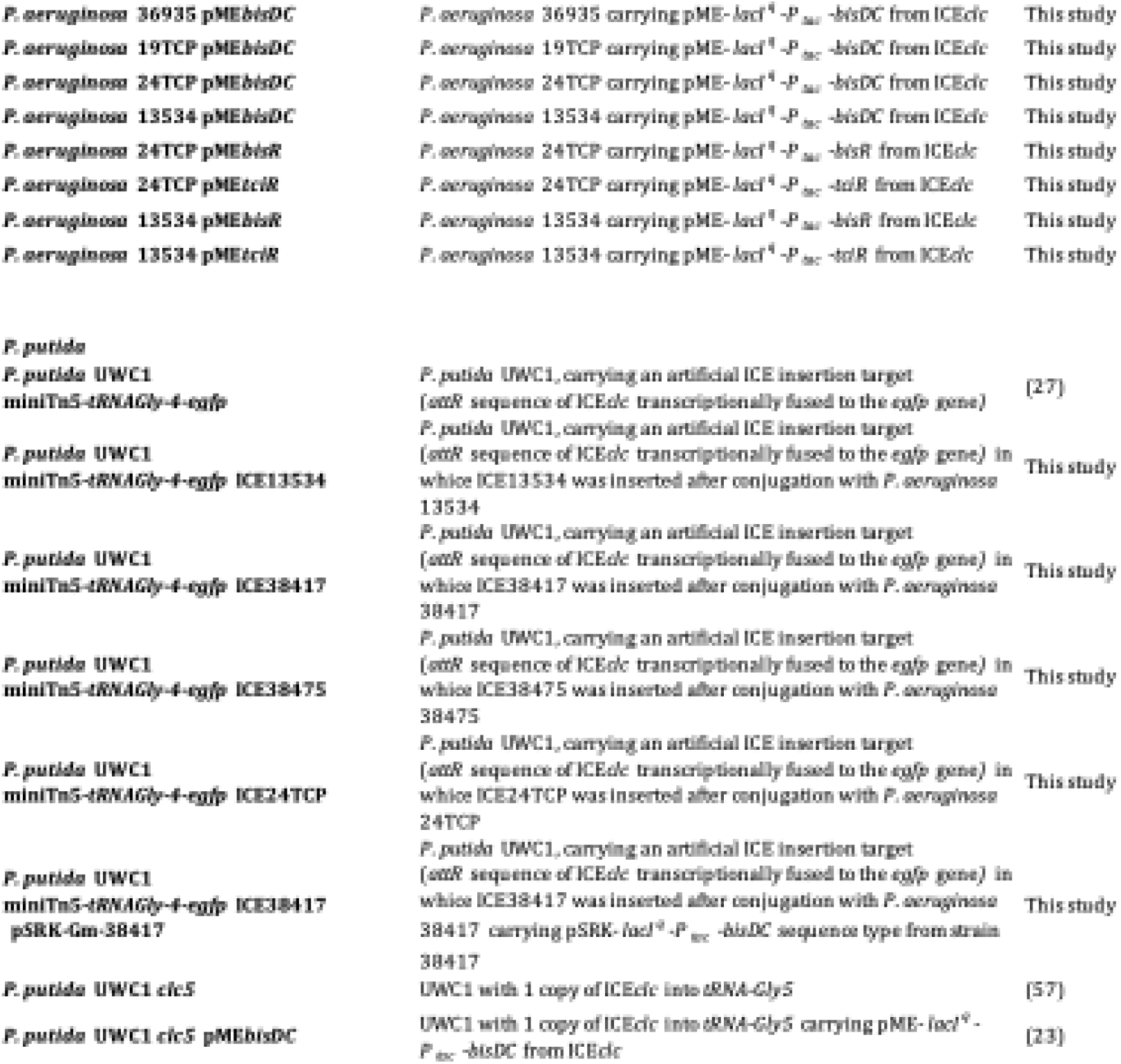
Strains used in this study.

**Supplementary Table 2:** Plasmids used in this study.

| Plasmids | Description | Reference |
| --- | --- | --- |
| pMEbisDC | TcR, *lacIq*-P<sub>lac</sub>-bisDC (allows IPTG-controlled expression of BisDC) | (23) |
| pMEbisR | TcR, *lacIq*-P<sub>lac</sub>-bisR (allows IPTG-controlled expression of BisR) | (23) |
| pMEtcIR | TcR, *lacIq*-P<sub>lac</sub>-tcIR (allows IPTG-controlled expression of TcIR) | (23) |
| pSRK | Broad-host-range expression vector containing lac promoter and *lacIq*, *lacZa*<sup>+</sup>, and TcR or GmR | (58) |
| pSRK-Gm-38417 | pSRK with GmR and *lacIq*-P<sub>lac</sub>-bisDC from strain 38417 | This study |
| pSRK-Gm-35866 | pSRK with GmR and *lacIq*-P<sub>lac</sub>-bisDC from strain 35866 | This study |
| pSRK-Gm-37327 | pSRK with GmR and *lacIq*-P<sub>lac</sub>-bisDC from strain 37327 | This study |
| pSRK-Gm-19TCP | pSRK with GmR and *lacIq*-P<sub>lac</sub>-bisDC from strain 19TCP | This study |

**Supplementary Table 3:**
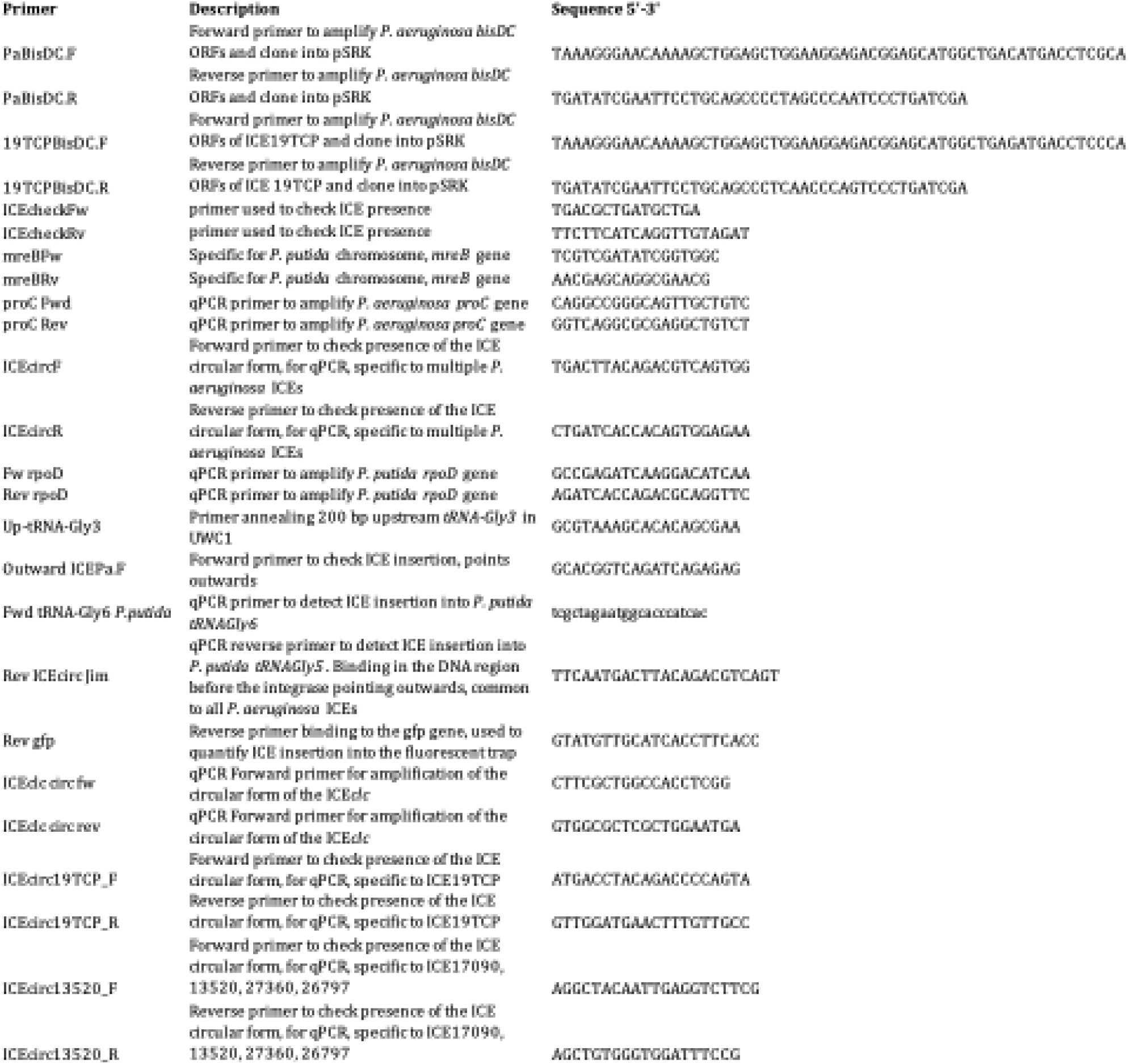
Oligonucleotides used in the study.

**Supplementary Table 4:**
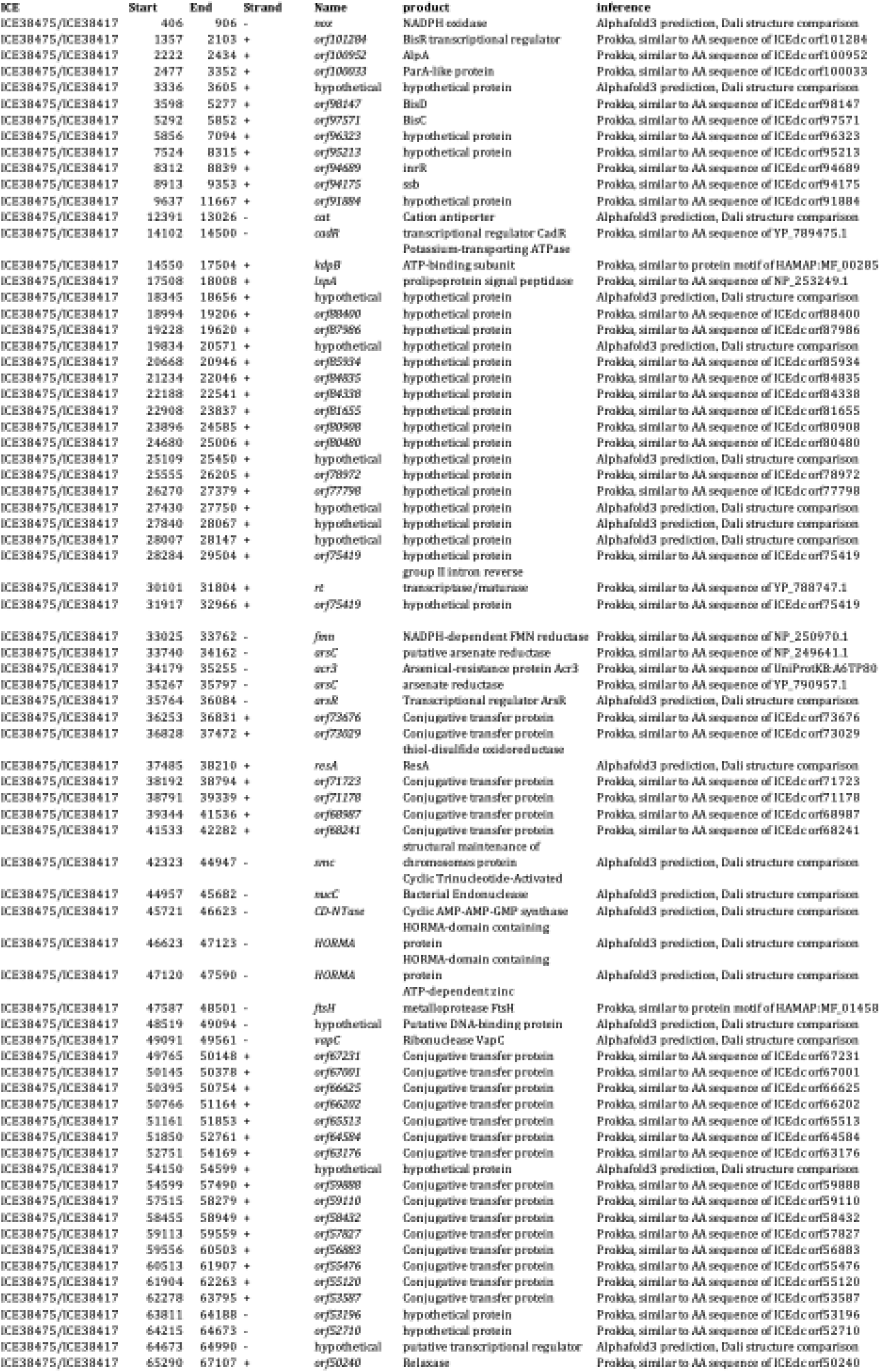
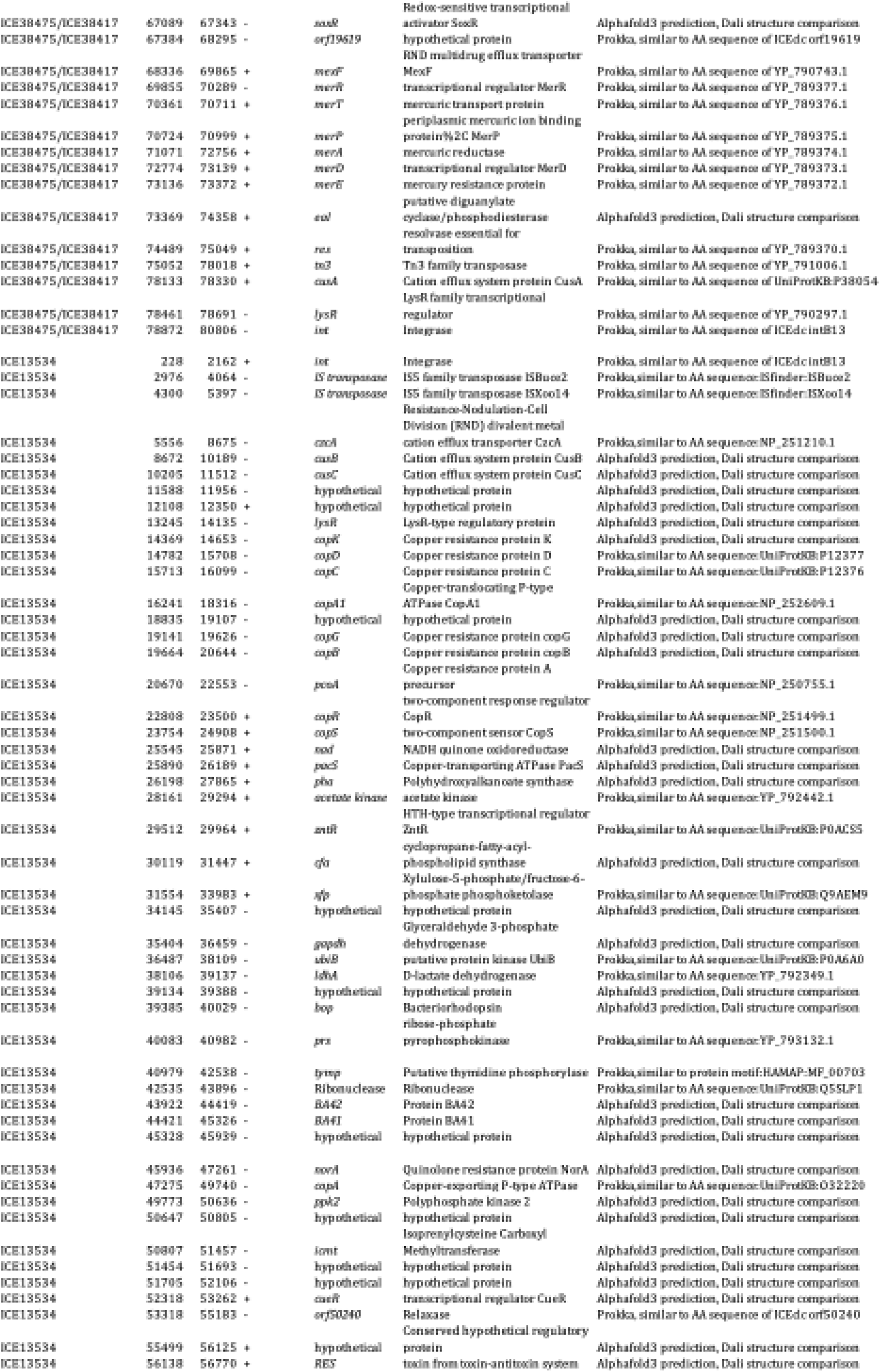
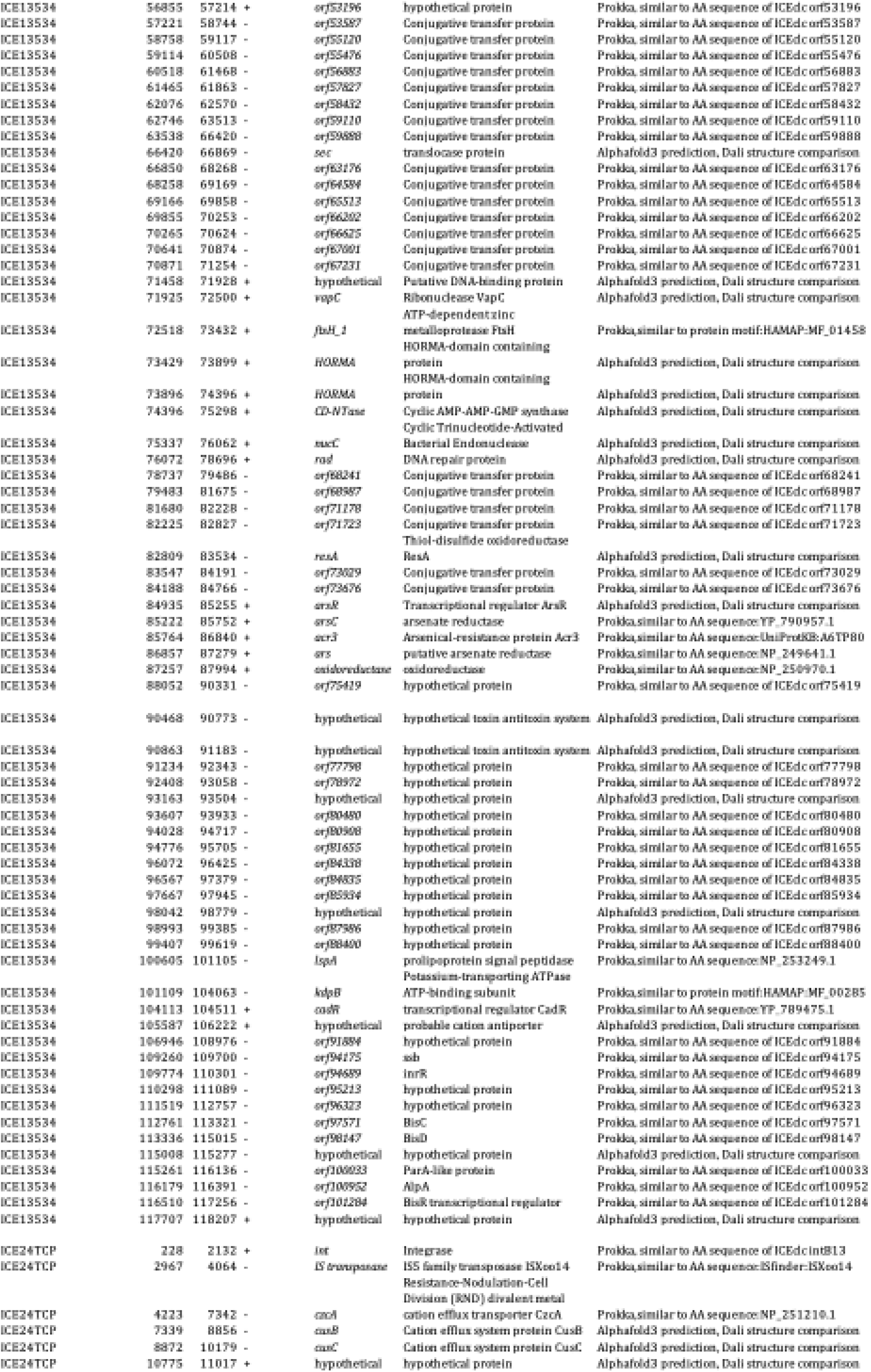
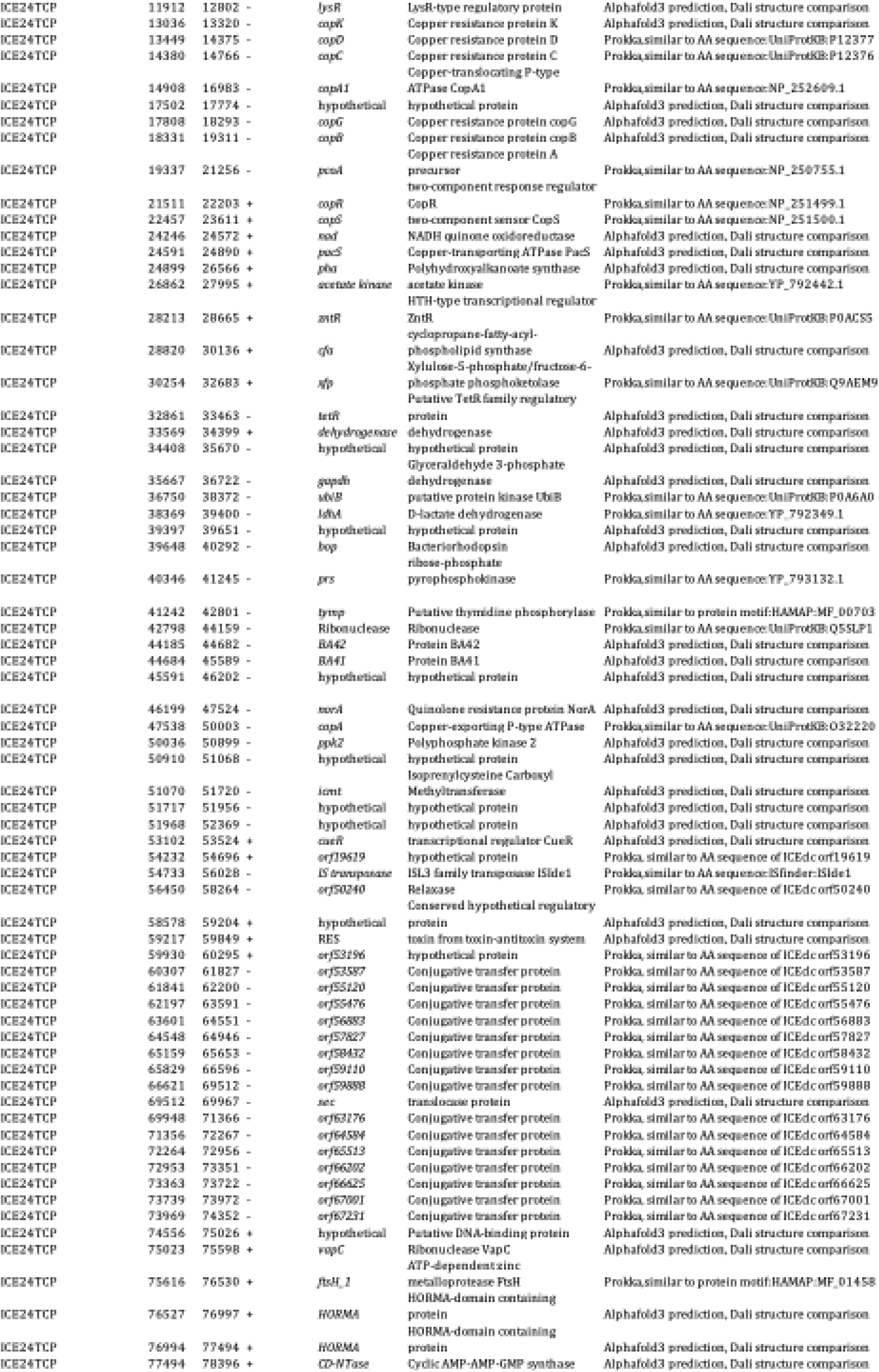
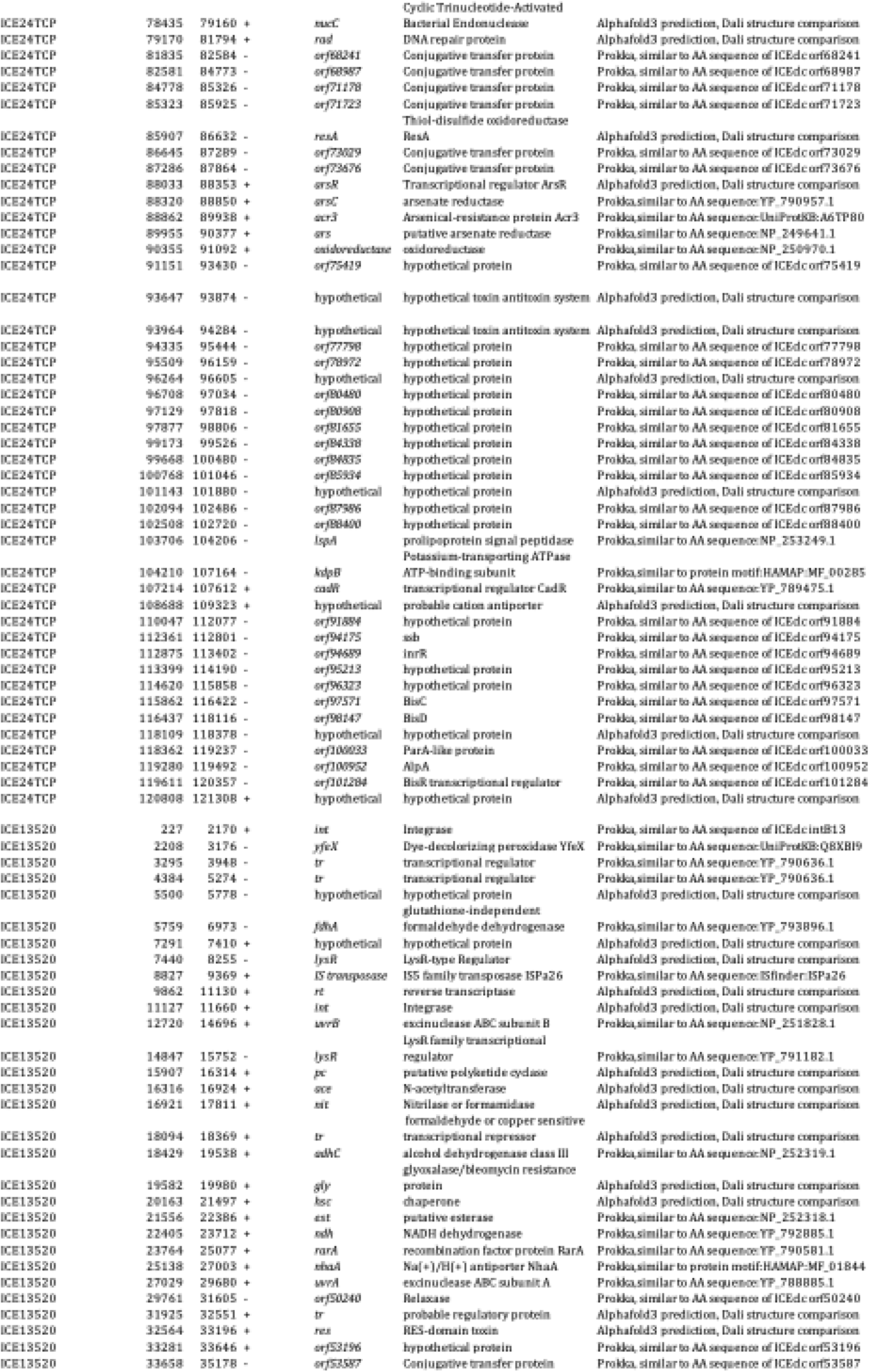
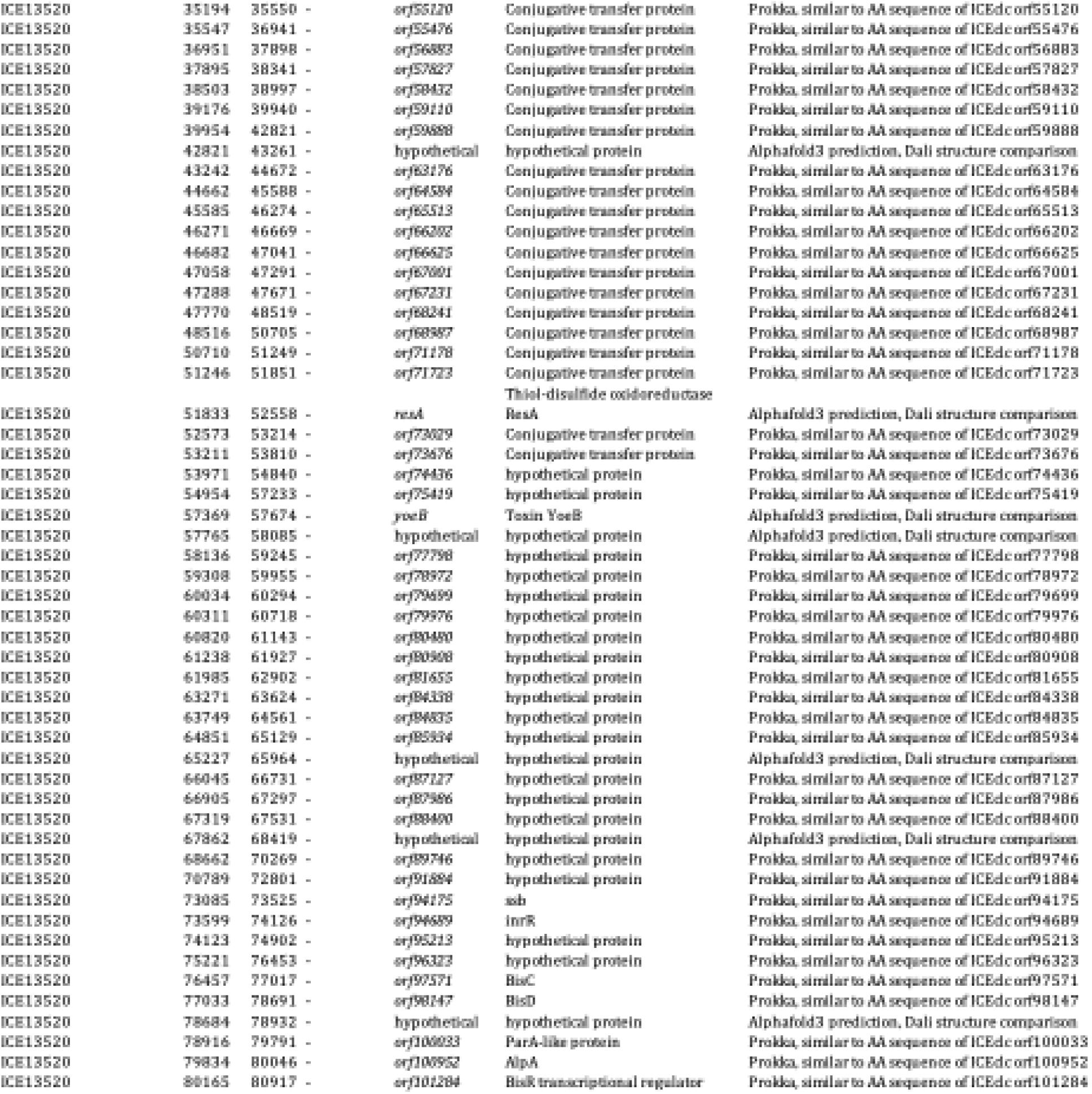
ICE annotation.

**Supplementary Table 5:** MIC of heavy metals in *P. putida* carrying ICEs from *P. aeruginosa*.

| Strain | Putative resistance | Number of ORFs | Hg(II) MIC in LB (μM) | Cu(II) MIC in LB (mg/L) | As(III) MIC in LB (mM) | Ciprofloxacin MIC on MH agar (mg/L) |
| --- | --- | --- | --- | --- | --- | --- |
| UWC1 | - | - | 10 | > 250 | > 15 | 0.047 |
| UWC1 ICE13534 | As | 4 | 10 | > 250 | > 15 | 0.047 |
|  | Cu | 13 |  |  |  |  |
|  | quinolones | 1 |  |  |  |  |
|  | Hg | 1 |  |  |  |  |
| UWC1 ICE38417 | As | 4 | > 40 | > 250 | > 15 | 0.016 |
|  | Hg | 6 |  |  |  |  |
| UWC1 ICE24TCP | As | 4 | 10 | > 250 | > 15 | 0.064 |
|  | Cu | 13 |  |  |  |  |
|  | quinolones | 1 |  |  |  |  |
|  | Hg | 1 |  |  |  |  |
| UWC1 ICE38475 | As | 4 | > 40 | > 250 | > 15 | 0.047 |
|  | Hg | 6 |  |  |  |  |

**Supplementary Table 6:** MIC of *P. aeruginosa* 13534 of different antibiotics. Susceptible (S), Susceptible, increased exposure (I), Resistant (R).

| Antibiotic | MIC (mg/L) | EUCAST Classification |
| --- | --- | --- |
| ceftazidime | 1 | I |
| cefepime | 2 | I |
| ciprofloxacin | 1.5-2 | R |
| meropenem | 0.094 | S |
| imipenem | 1.5 | I |
| ceftazidime avibactam | 0.5-0.75 | S |
| ceftolozane tazobactam | 0.75-1.5 | S |
| ceftaroline | 6 - 8 | NA |
| colistine | 2 | S |
| amikacine | 6 | S |
| tobramycine | 1.5 | S |

**Supplementary Table 7:**
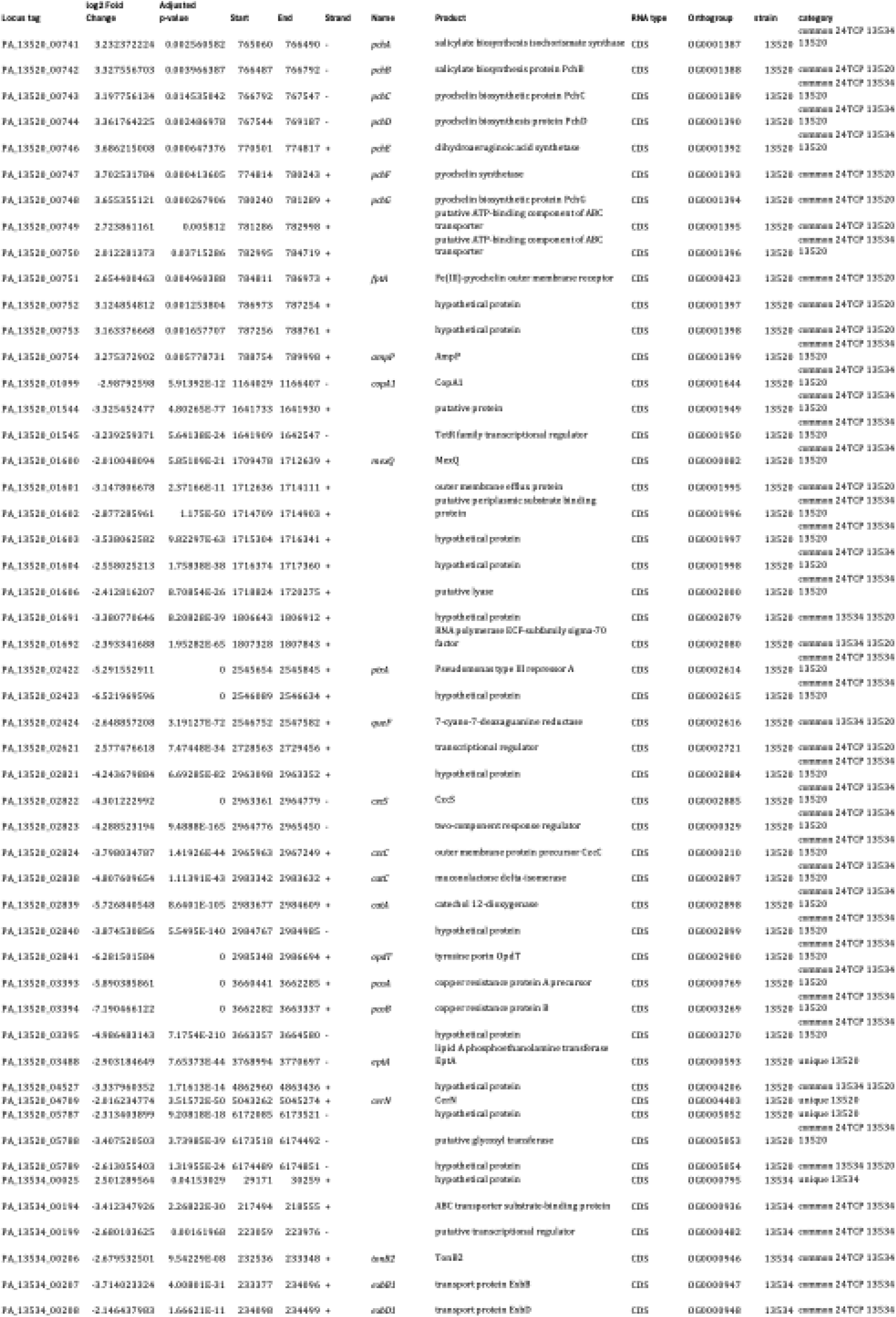
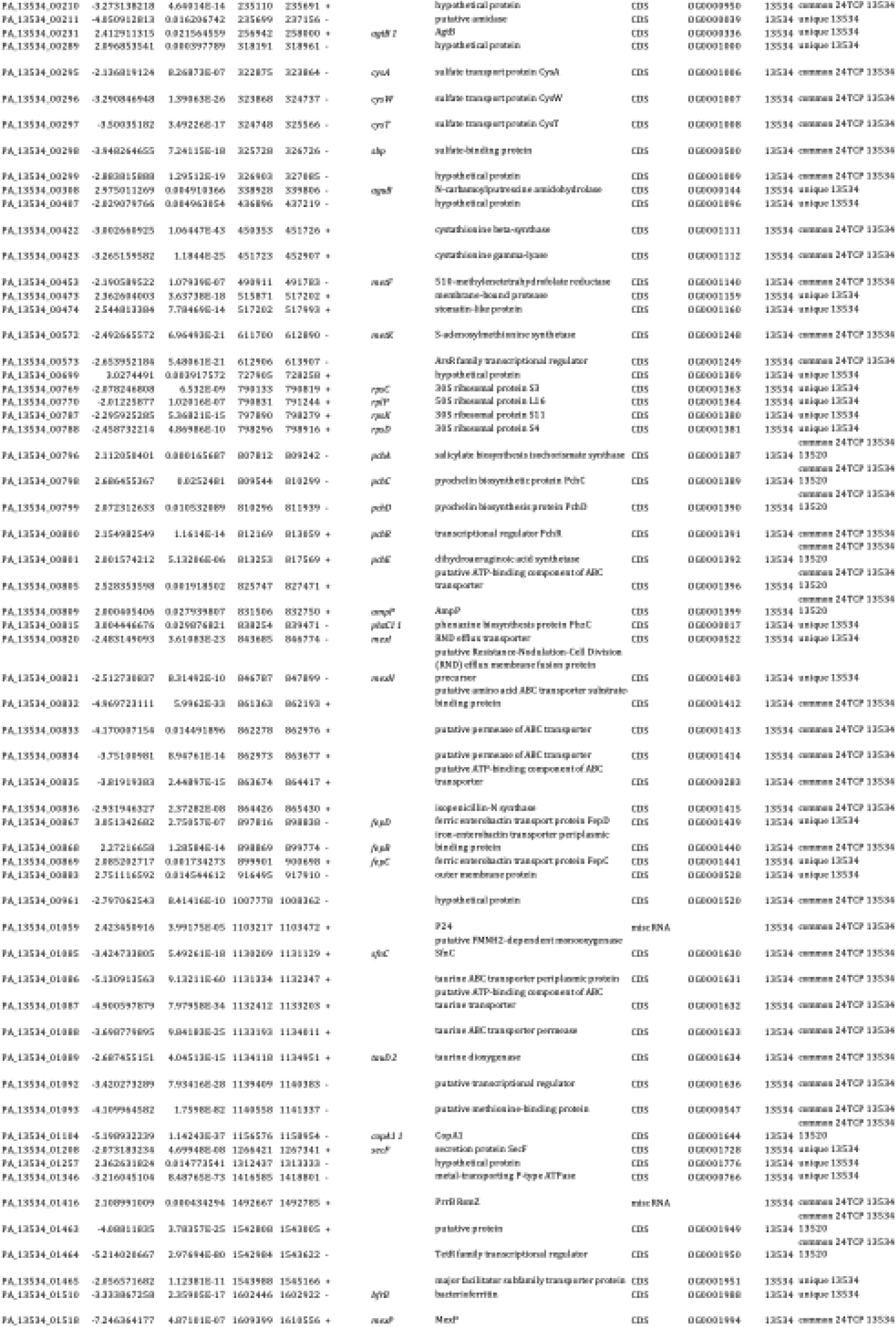
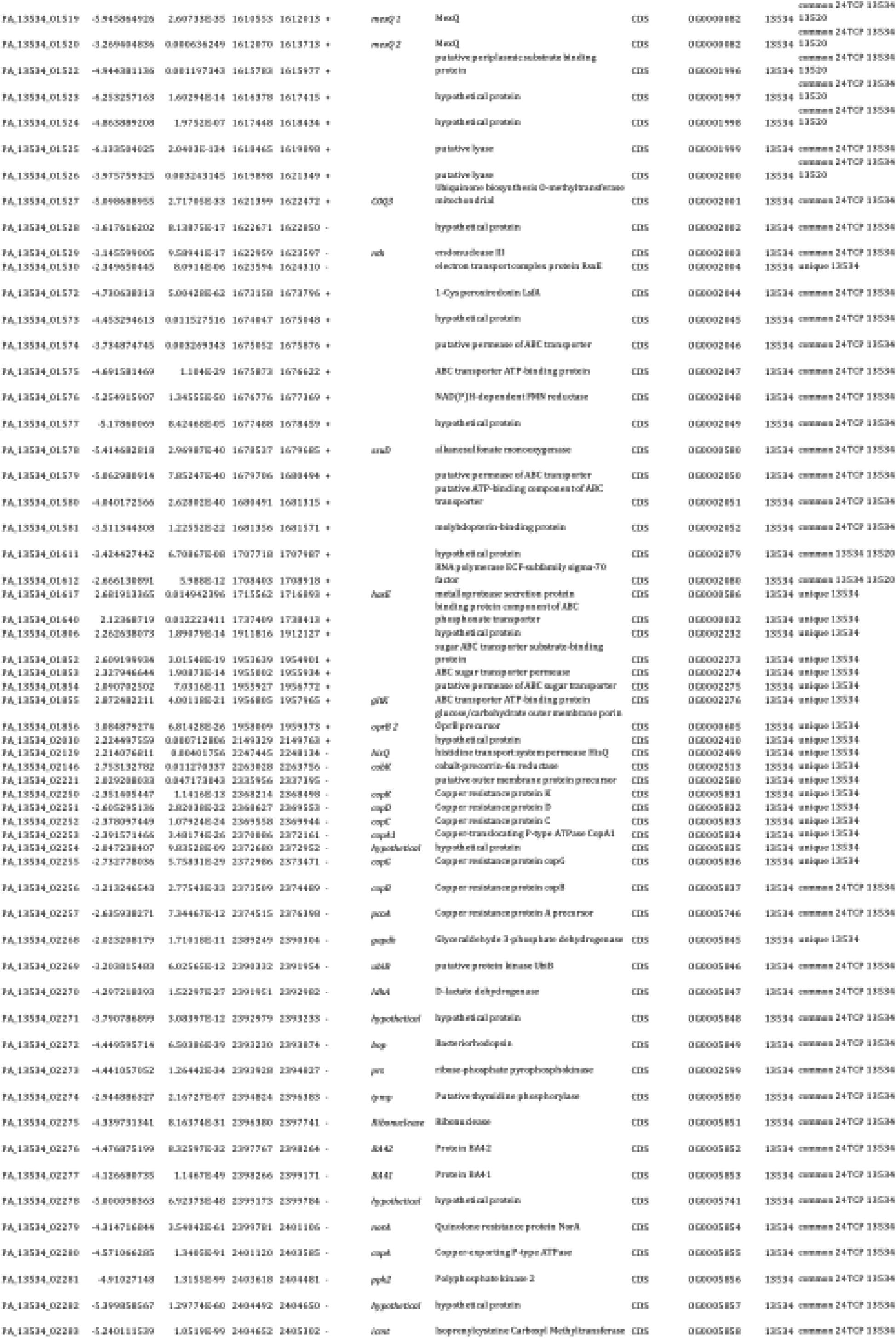
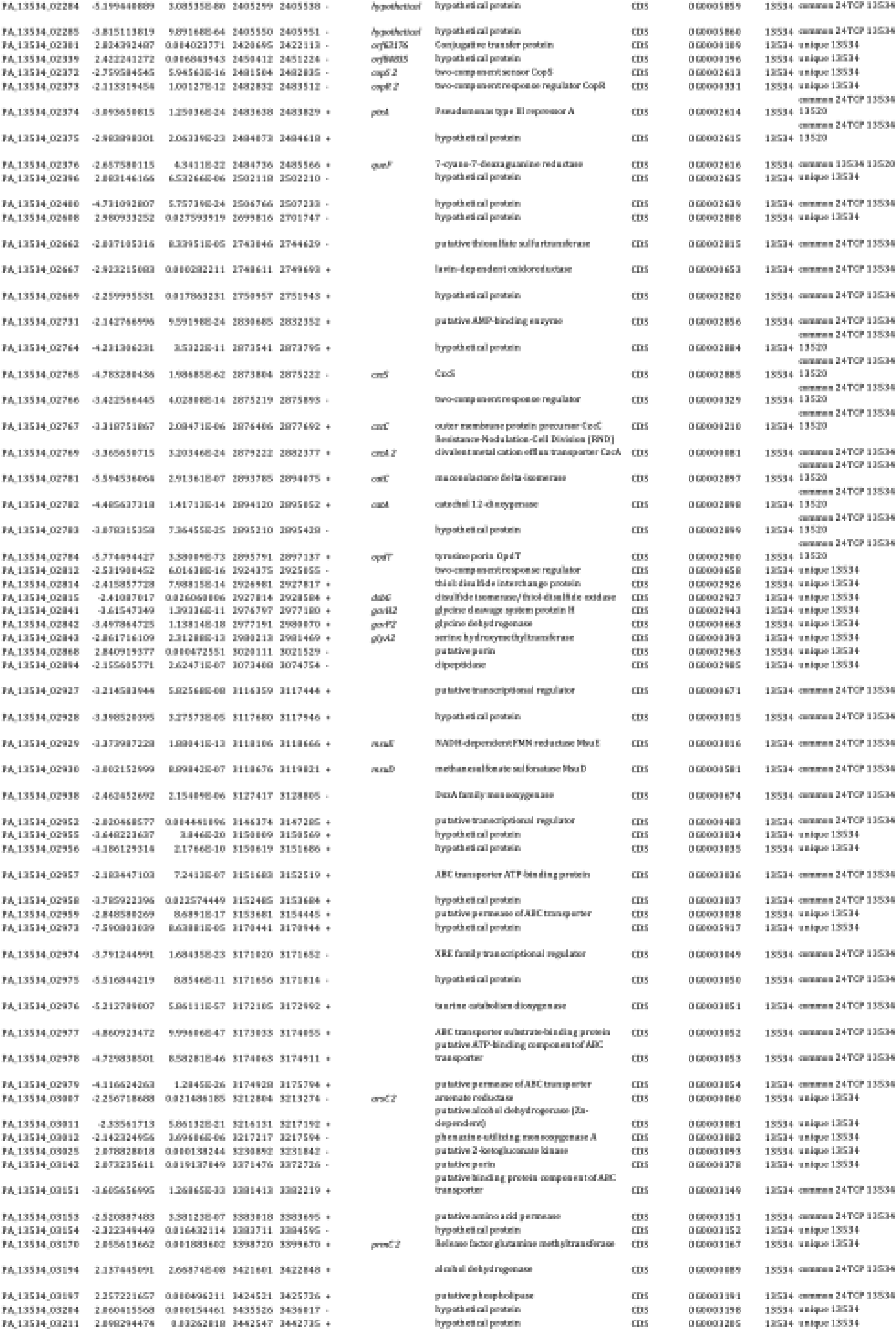
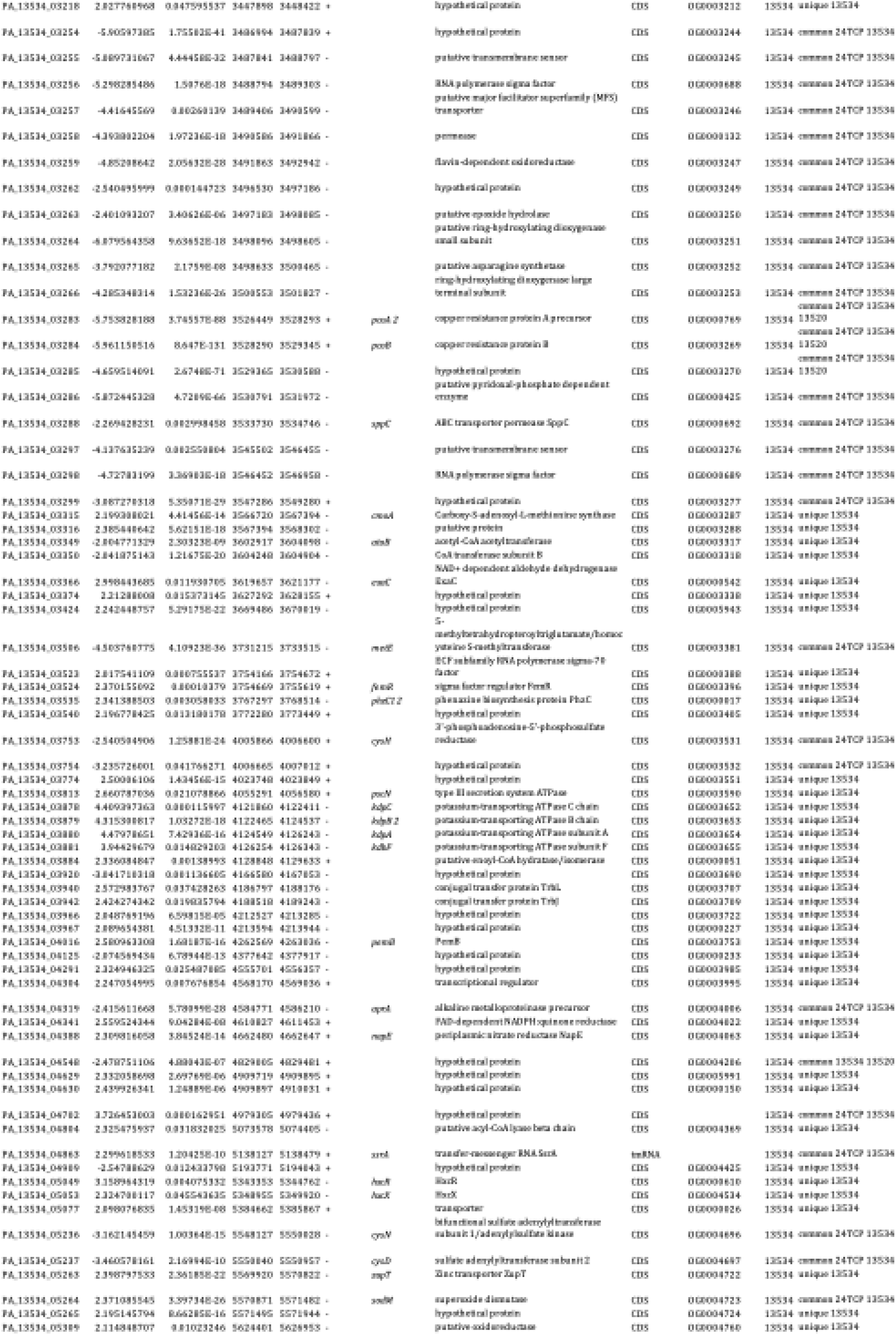
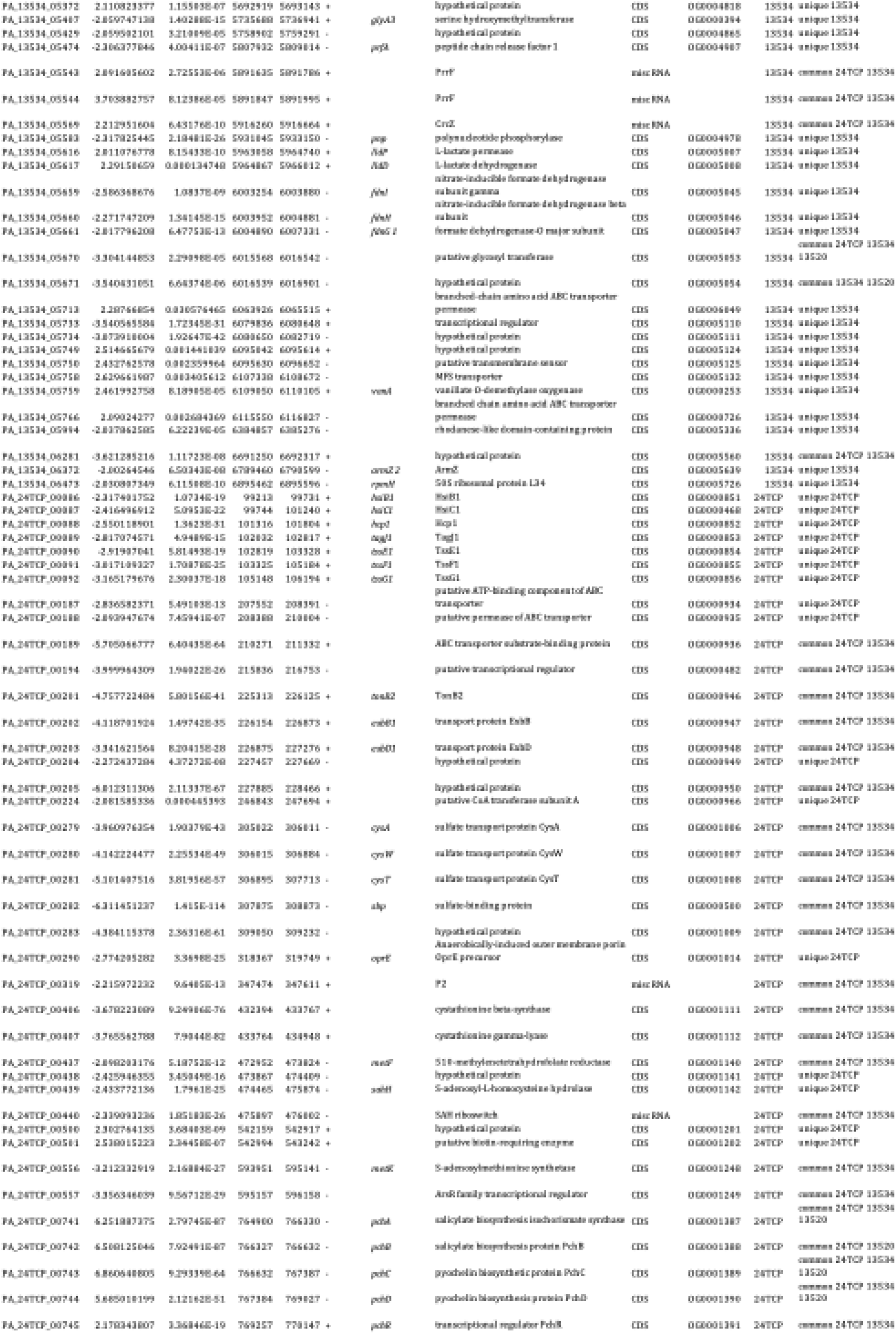
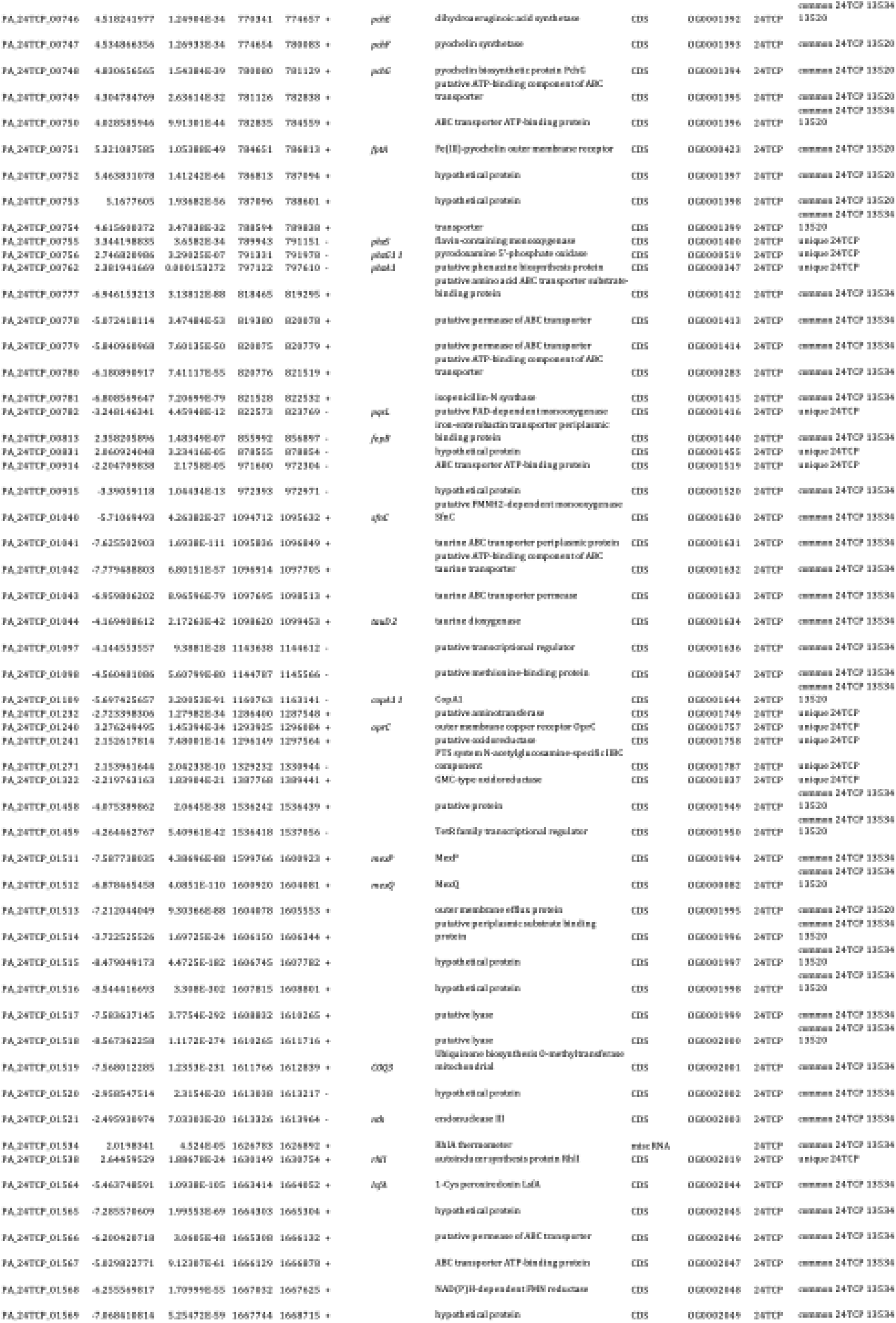
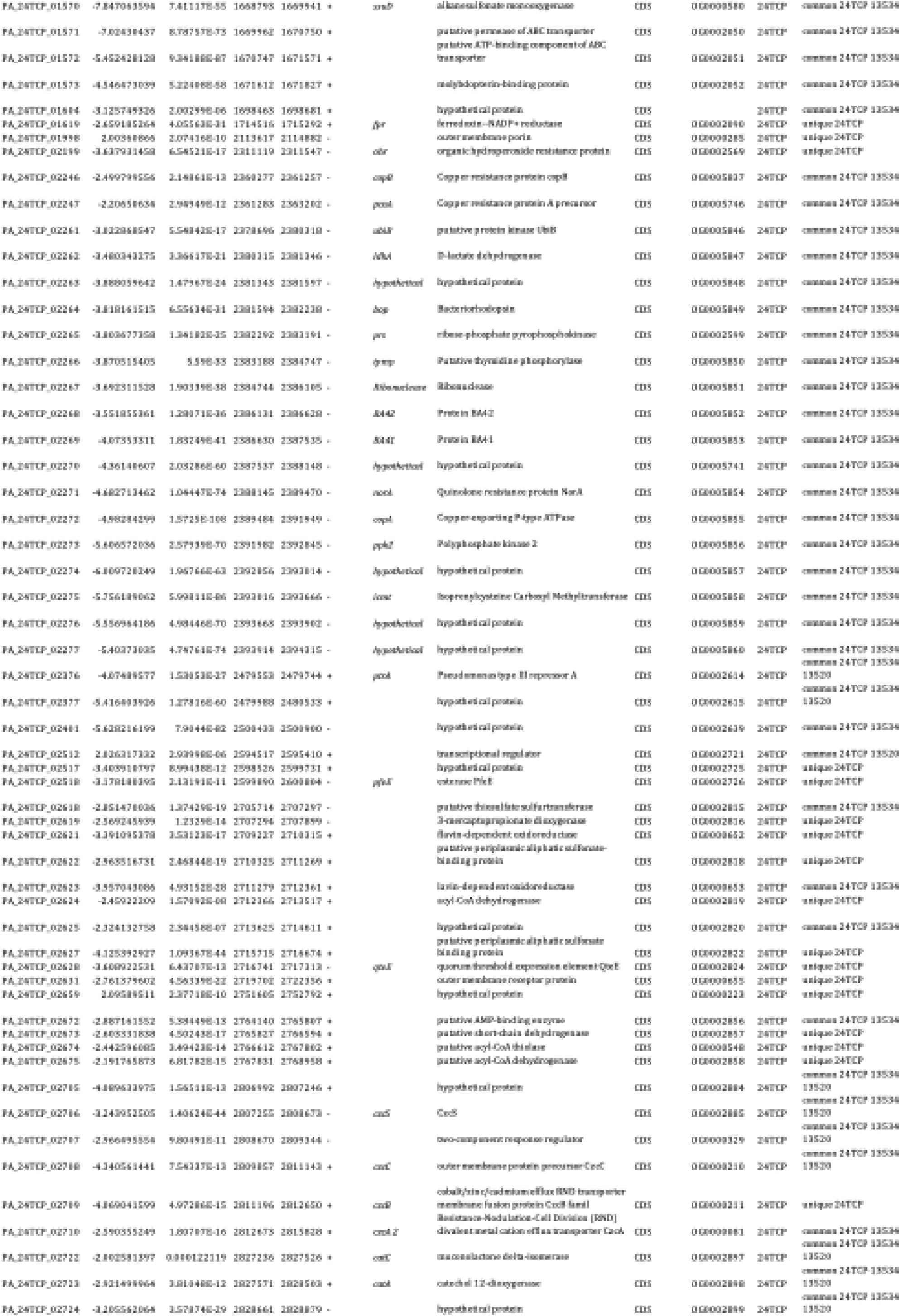
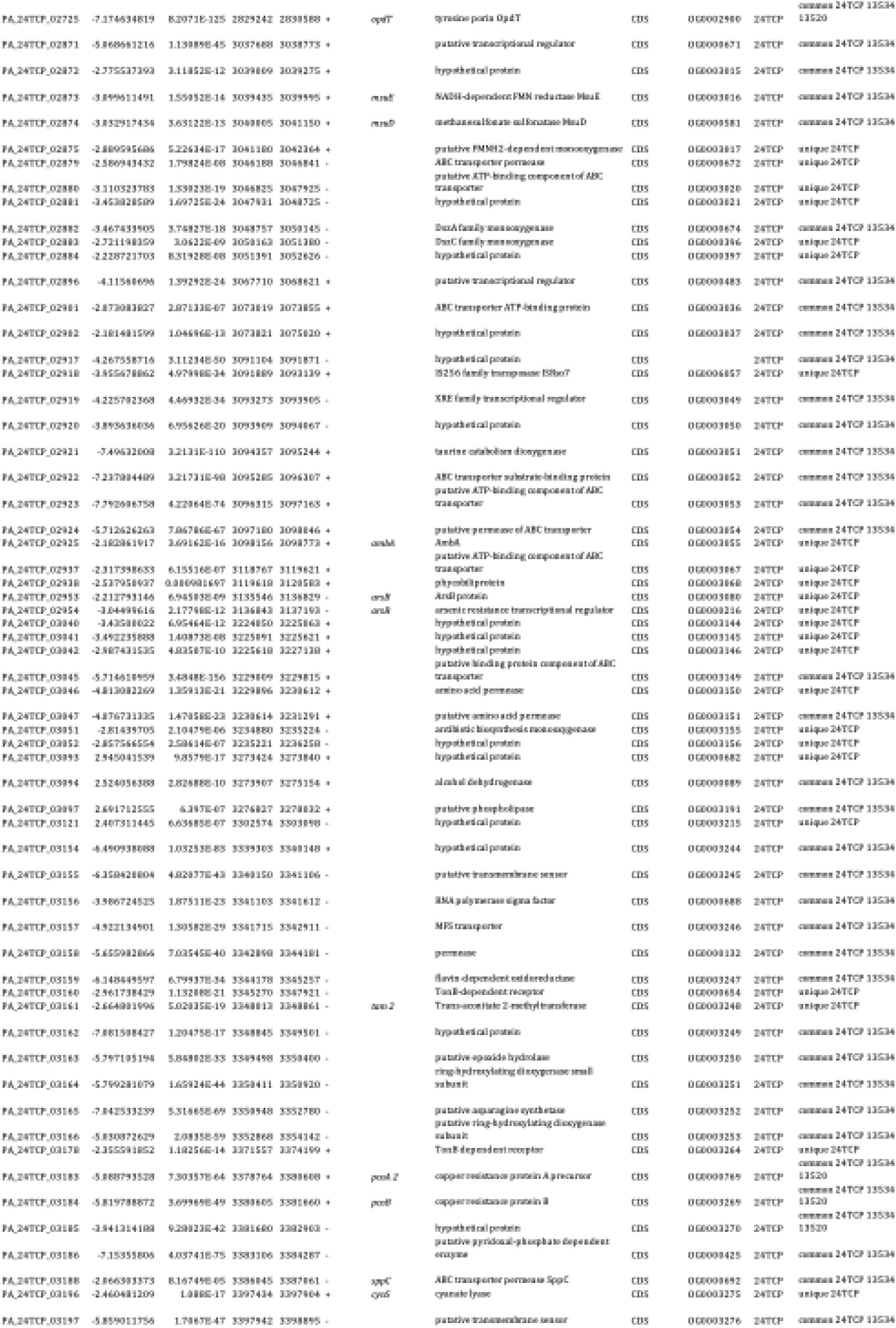
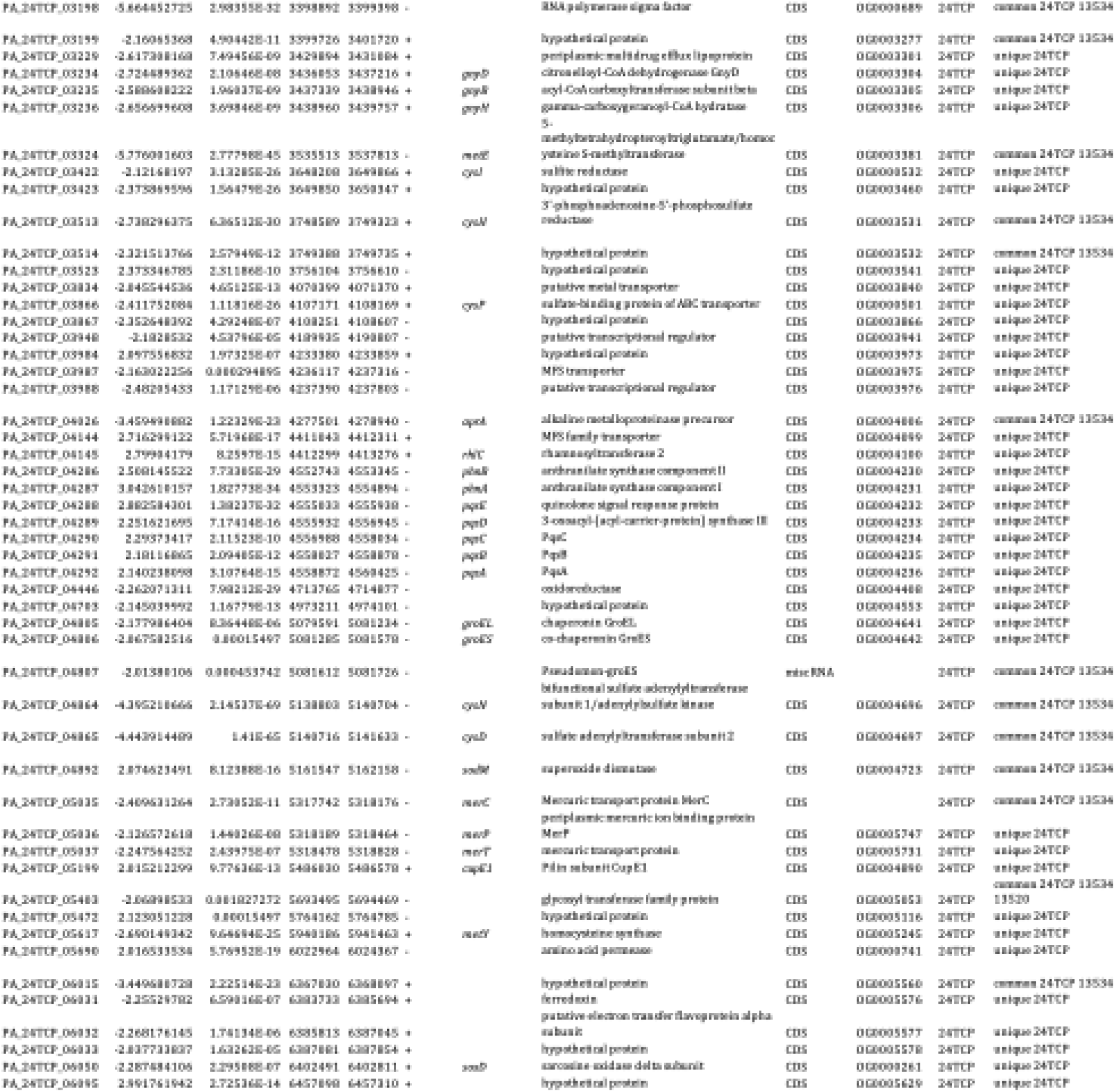
Differentially expressed genes in *P. aeruginosa* 13534, 24TCP, 13520 growing on LB vs LB + 50 mg/L Cu.

